# Evolution of gene-rich germline restricted chromosomes in black-winged fungus gnats through introgression (Diptera: Sciaridae)

**DOI:** 10.1101/2021.02.08.430288

**Authors:** Christina N. Hodson, Kamil S. Jaron, Susan Gerbi, Laura Ross

## Abstract

Germline restricted DNA has evolved in diverse animal taxa, and is found in several vertebrate clades, nematodes, and flies. In these lineages, either portions of chromosomes or entire chromosomes are eliminated from somatic cells early in development, restricting portions of the genome to the germline. Little is known about why germline restricted DNA has evolved, especially in flies, in which three diverse families, Chironomidae, Cecidomyiidae, and Sciaridae exhibit germline restricted chromosomes (GRCs). We conducted a genomic analysis of germline restricted chromosomes in the fungus gnat *Bradysia* (*Sciara*) *coprophila* (Diptera: Sciaridae), which carries two large germline restricted “L” chromosomes. We sequenced and assembled the genome of *B. coprophila*, and used differences in sequence coverage and k-mer frequency between somatic and germ tissues to identify GRC sequence and compare it to the other chromosomes in the genome. We found that the GRCs in *B. coprophila* are large, gene-rich, and have many genes with paralogs on other chromosomes in the genome. We also found that the GRC genes are extraordinarily divergent from their paralogs, and have sequence similarity to another Dipteran family (Cecidomyiidae) in phylogenetic analyses, suggesting that these chromosomes have arisen in Sciaridae through introgression from a related lineage. These results suggest that the GRCs may have evolved through an ancient hybridization event, raising questions about how this may have occurred, how these chromosomes became restricted to the germline after introgression, and why they were retained over time.

## Introduction

An underlying tenet of heredity is that all cells within an organism have the same genomic sequence. However, there are a surprising number of exceptions to this rule. For instance, Boveri [1] noted in *Ascaris* nematodes that fragments of chromosomes were eliminated from somatic cells early in development, showing that in some cases germline/ soma differentiation involves changes in the genomic composition of cells as well as regulatory changes. In addition to the loss of chromosomal fragments (referred to as “chromatin diminution”), another type of germline specialization involves the elimination of whole chromosomes from somatic cells. A phenomenon we believe this was first noted in the Dipteran gnat *Bradysia* (*Sciara*) *coprophila* [2]. Both chromatin diminution and chromosome elimination are examples of programmed DNA elimination, which occurs in a developmentally regulated manner across a broad evolutionary range from ciliates to mammals, including more than 100 species from nine major taxonomic groups [3]. Programmed DNA elimination is not a rare phenomenon, yet remains poorly understood. Recently, however, genomic studies in several species are beginning to address questions regarding their function and evolution.

Many examples of programmed DNA elimination involve regulated DNA elimination from somatic cells so that portions of the genome are restricted to the germline [3]. Germline restricted DNA, involving either portions of chromosomes (chromatin diminution) or entire chromosomes (chromosome elimination) have evolved repeatedly and are found in lampreys and hagfish (the most basal vertebrates), songbirds, nematodes, and flies [1,4–7]. Recent genomic work on lampreys and nematodes (with chromatin diminution) and songbirds (with chromosome elimination) have found that the germline restricted portions of the genome often carry protein coding genes involved in germ tissue maturation and function [8–11]. Therefore, a leading hypothesis is that germline restricted DNA may help resolve intralocus conflict between the germline and somatic cells [10,12]. However, although chromatin diminution and chromosome elimination have similar consequences, the initial evolution of these systems probably differs, as the mechanism of elimination is substantially different in these two systems.

In species with chromosome elimination, entire chromosomes are exclusively found in the germline: the germline restricted chromosomes (GRCs). Little is known about how these chromosomes arise and how they are related to the rest of the genome. One hypothesis is that they originate from B chromosomes [13], which are accessory non-essential chromosomes that are widespread in eukaryotes [14]. GRCs are similar to B chromosomes in that they are chromosomes in addition to the core genome (i.e. the chromosomes which are found in the somatic cells as well as the germ cells), with greater variation in presence/number of chromosomes than the core chromosome set. However, while B chromosomes are non-essential, recent genomic work in songbirds suggests that GRCs likely play an important, and perhaps fundamental role in zebra finches [10] and are evolutionarily conserved across songbirds [15]. Furthermore, there is no clear evidence that GRCs spread through drive and therefore unlike B chromosomes most likely persist due to their functional importance, rather than as reproductive parasites. So while it is still possible that GRCs originated from B chromosomes and were subsequently “domesticated”, alternative explanations for their origin cannot be excluded. Especially as the origins of the GRCs have so far only focused on their single origin among birds. Here we focus on a different origin of GRCs; their evolution and origin in flies (Diptera).

GRCs are found in three dipteran families: the “K” chromosomes of non-biting midges (Chironomidae), the “E” chromosomes of gall gnats (Cecidomyiidae), and the “L” chromosomes of black winged fungus gnats (Sciaridae) [4,16,17]. Each instance appears to have an independent origin, as GRCs show different properties in each lineage, and the three families are not sister clades [18,19]. While the evolutionary origins of these chromosomes remain obscure, GRCs are expected to have some function relating to reproduction, otherwise, they likely would not have been retained over time. The origin and evolution of GRCs in Sciaridae and Cecidomyiidae are particularly intriguing, as these families are relatively closely related, both belonging to the infraorder Bibionomorpha (although they are not sister clades, [19]). Therefore, understanding how GRCs arose in these two lineages and what factors led to their evolution can provide a foundation from which we can answer many questions. For instance, we can start to unravel why GRCs arose in some Bibionomorpha families but not others, and compare the gene content and expression of GRC genes in two relatively closely related families.

Although both Sciaridae and Cecidomyiidae carry GRCs, the characteristics of these chromosomes differ between the two families, with Sciaridae carrying few (up to 4) large GRCs, and Cecidomyiidae carrying many (between 16 and 67) small GRCs (reviewed in [18,20]). Therefore, theories for how GRCs arose differ between the two lineages. In Cecidomyiidae, the GRCs show some similarities in appearance to the core genome, and so it was originally proposed that they evolved through whole genome duplications followed by restriction of the duplicated chromosomes to the germline [21,22]. However, this idea remains controversial and lacks empirical support. In Sciaridae, however, a comprehensive theory for the evolution of GRCs suggests that the GRCs evolved from the X chromosome in a series of conflicts between different parts of the genome [23]. This theory suggests that the evolution of GRCs is closely intertwined to the unusual genetic system found in this lineage. Sciaridae displays a non-Mendelian chromosome inheritance system known as paternal genome elimination [16,24] and has an XO sex chromosome system. In species with paternal genome elimination, meiosis in males is unconventional such that males only transmit chromosomes that they inherit from their mother to their offspring, while paternal chromosomes are eliminated in male meiosis. In addition, in *B. coprophila* male meiosis is also unusual in that all GRCs present (normally two) are transmitted to offspring through sperm, and there is an unusual X chromosome nondisjunction event such that two copies of the X chromosome are transmitted through sperm, resulting in males transmitting two GRCs, two X chromosomes, and a haploid set of autosomes through sperm (**Fig 1**). Furthermore, the sex determining X chromosomes in Sciaridae are not inherited from the parents, instead the sex is determined by the number of X chromosomes eliminated from somatic cells early in development [2,25]. Sex chromosome elimination occurs in early in development, when the X chromosome(s) that will be eliminated are left on the metaphase plate and not incorporated into daughter nuclei. GRCs are eliminated from somatic cells in a similar way, with the exception that GRC elimination occurs slightly earlier in development than X chromosome elimination [2] (**Fig 1**; see **Supplementary Text 1** for additional information).

**Fig 1.**
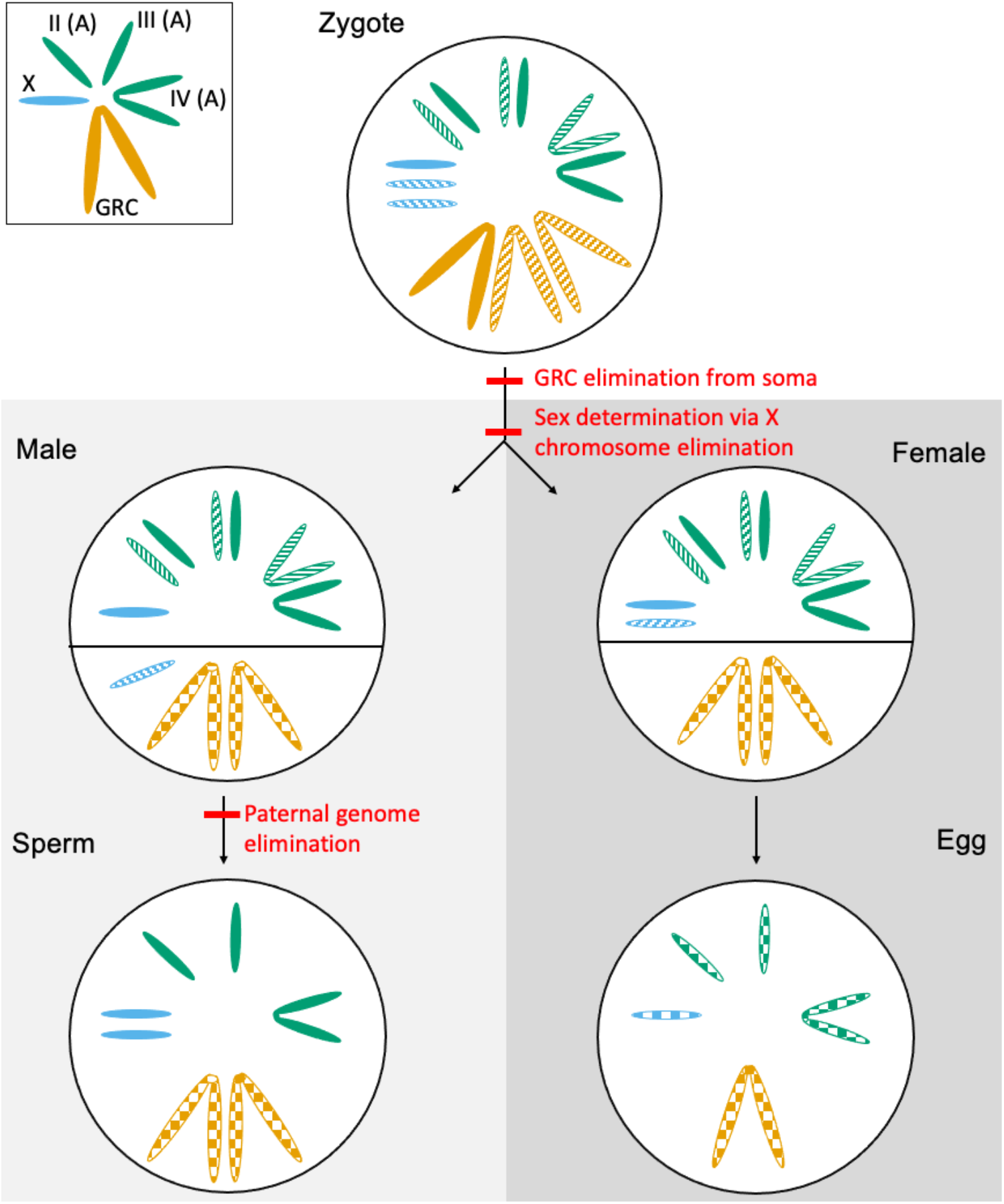
Chromosome dynamics during *B. coprophila* development. *Bradysia coprophila* has three autosomes (A, green), an XO sex determination system (X chromosome shown in blue), and germline restricted chromosomes (GRC, shown in orange). paternal origin chromosomes = dashed, maternal origin chromosomes = solid, either maternal or paternal origin chromosomes = chequered. Chromosomes below the solid line in males and females are additional chromosomes present in the germ tissue but eliminated from somatic tissue. *Bradysia coprophila* GRCs are eliminated from somatic cells early in development and X chromosome elimination (always paternally inherited X chromosomes) are eliminated early in development from somatic cells to determine sex. Males also undergo paternal genome elimination such that (apart from the GRCs) only maternally inherited chromosomes are transmitted through the sperm (including two copies of the maternally derived X chromosome due to a non-disjunction event in meiosis). Chromosome sizes and shapes approximated from [16].

Haig’s theory [23] suggests that paternal genome elimination and X chromosome elimination as a means of sex determination evolved at the base of the Sciaridae. Following this, GRCs evolved from the paternally derived X chromosome in males as a means to escape elimination through paternal genome elimination. This was followed by restriction of this chromosome to the germline as X chromosome polyploidy in the somatic cells might be detrimental. Although, there has been no attempt to validate this theory in Sciaridae, it contains some testable predictions. For instance, following this theory [23], we would expect that the GRCs, if they were derived from the X chromosome, would exhibit some homology to this chromosome, and that the GRCs would be of relatively recent origin, originating within the Sciaridae. Interestingly, Cecidomyiidae species also exhibit paternal genome elimination and X chromosome elimination as a means of sex determination [18,26]. However, if Haig’s theory is correct GRCs, paternal genome elimination, and X chromosome elimination as a means of sex determination evolved independently in these two clades. There is recent evidence suggesting that the X chromosomes in Cecidomyiidae and Sciaridae are not related [27], but besides this, how the reproduction systems in both the Cecidomyiidae and Sciaridae evolved remains a mystery and very little empirical work has been done on this topic in either clade.

We conduct the first genomic analysis of GRCs in Diptera, with the goal of exploring the origin, evolution, and structure of GRCs in Sciaridae. GRCs in Sciaridae are historically referred to as L chromosomes, however we refer to them as GRCs in this paper to more easily facilitate comparison with GRCs in other lineages. We sequence germline and somatic tissue from *B. coprophila* and identify GRC scaffolds in a genome assembly generated from both tissue types by comparing coverage levels and k-mer distributions between the two sequence types (with the idea that GRC sequences will be present in the germline but not in the soma). We were able to unambiguously identify GRC scaffolds and perform downstream analyses to compare the gene-content between GRCs, autosomes and the X chromosome in *B. coprophila*.

We find that the two GRCs are gene-rich and carry many paralogs to the core genome. Contrary to Haig’s theory, we do not find significant homology of GRCs to the X chromosome, rather, we find GRC paralogs throughout the genome with high levels of divergence from the GRC copy. Remarkably, phylogenomic analyses suggest that the GRCs in *B. coprophila* arose through introgression from distantly related cecidomyiids. This clade also carries GRCs but these were previously assumed to have originated independently. How this ancient introgression occurred, why these chromosomes were retained, and how they became restricted to the germline are intriguing questions raised from our results. This study provides a foundation for the study of GRCs in Sciaridae, an understudied lineage with regards to GRCs, with great potential given the rich body of molecular and cytological research in Sciaridae researched for nearly a century [20,28,29]. Furthermore, fungus gnats are cosmopolitan species easy to rear in the laboratory, allowing for future studies of function and diversity of GRCs in the family. This study also adds to the recent genomic studies on germline restricted DNA in animals, suggesting that germline restricted DNA often contains numerous protein coding genes.

## Results and discussion

One consequence of the unconventional genetic system in *B. coprophila* is that male somatic and germ cells have a different chromosome constitution. They differ in the presence of germline restricted chromosomes, but also in the frequency of X chromosomes (two are present in germ cells, but only one is present in somatic cells) (**Fig 2A**). We used these differences in chromosome constitution to identify the GRC chromosomes in *B. coprophila*, and also to differentiate the X chromosome from autosomes. We sequenced adult male germ and somatic tissue and generated a genome assembly from both the germ and somatic sequence libraries. (See Methods and Supplementary Text 2 for assembly information). The genome assembly is of a comparable size to flow cytometry estimates for the genome of *B. coprophila* [30] (**Table 1**).

**Table 1.** Size and gene content of autosomes (all three autosomes combined), X chromosome, and GRCs identified through k-mer and coverage differences between soma and germ tissue. Chromosome sizes are compared to flow cytometry estimates for *B. coprophila* [30] and gene number is compared to the reference genome assembly [31]. See **Supplementary Table 1** for assembly statistics.

|  | Size (Mb) |  | Gene Number |  |
| --- | --- | --- | --- | --- |
|  | Expected [30] | This study | Urban et al. [31] | This study |
| Whole genome | 362 | 398 | 23,117 | 41,418 |
| Autosomes | 225 | 162.4 | 18,254 | 17,802 |
| X | 49 | 52.9 | 4,863 | 4,277 |
| GRC | 88 | 154 | NA | 15,812 |
| Unclassified | NA | 20.2 |  | 3,527 |

**Fig 2.**
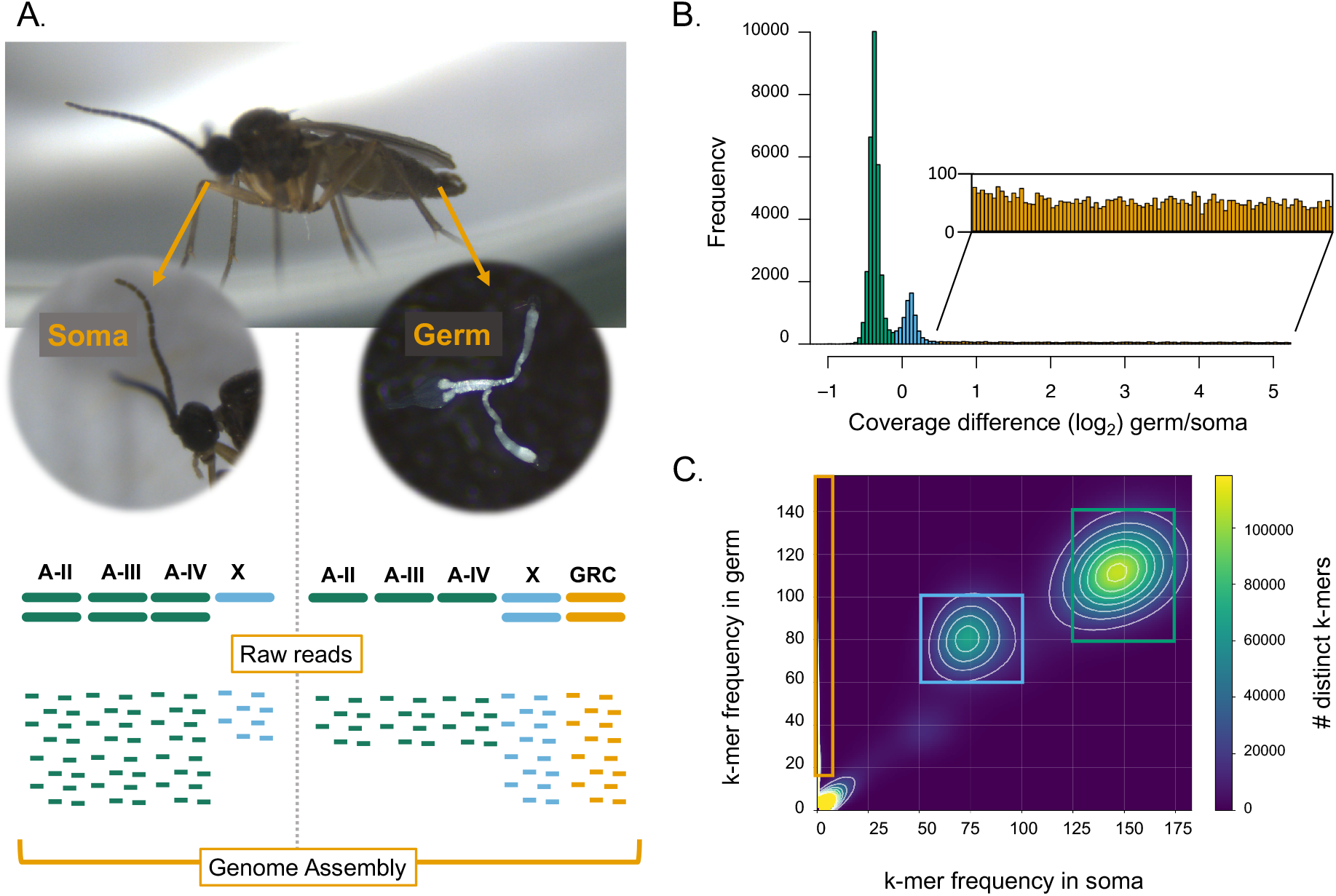
Sequencing and identification of GRC through comparison of germline and soma coverage. **A.**Schematic of sequencing approach for identifying GRC sequences in *Bradysia coprophila*.We isolated and sequenced somatic (head) and germ (testes with sperm) tissue. Somatic and germ tissue differ in the number of autosomes (A-II, A-III, and A-IV) (green), X chromosomes (blue), and GRCs (orange). We used differences in the chromosome constitution to isolate regions belonging to each chromosome type in a genome assembly made from short read sequences from both tissue types. **B**. Histogram of per scaffold log_2_ coverage differences between germ and somatic tissues. Green regions were assigned as autosomal, blue assigned as X-chromosome, and orange assigned as belonging to the GRCs (inset). **C**. Comparison of k-mer frequency differences between raw reads in the germ and soma libraries. K-mers were mapped to the genome assembly and scaffolds assigned based on which type of k-mers (GRC, X chromosome, or autosomal) mapped to the scaffold. Boxes show coverage of k-mers assigned as autosomal, X chromosome, and GRC.

### Bradysia coprophila GRCs are large and gene-rich

In order to identify the GRCs in our genome assembly, we utilized coverage differences and differences in k-mer profiles between the somatic and germ tissue sequencing libraries. We identified scaffolds that have a higher coverage in germ tissue than the somatic tissue (log_2_ germ/soma coverage difference > 0.5) (**Fig 2B**) and a high proportion (>80%) of GRC-specific 27-mers on the scaffold (**Fig 2C**, see **Materials and Methods** for details). We used a conservative approach, assigning GRC scaffolds only if both methods agreed on the assignment. Through this method we were also able to identify regions that belonged to the X chromosomes or autosomes. Through both the coverage and k-mer assignment of chromosomes, we identified 162.4 Mb of sequence as autosomal, 52.9 Mb of sequence that belong to the X chromosome, and 154 Mb of sequence that belong to the GRC (**Table 1**). The 20.2 Mb of sequence that we were unable to classify (**Table 1**) represent cases when the two methods (coverage and kmer-based) did not support the assignment with high confidence, indicating overall high agreement of the two approaches. With the exception of the GRC size, which is approximately double the size that we would expect given flow cytometry estimates of chromosome size in *B. coprophila* (Rasch, 2006), our chromosome size estimates are comparable to chromosome size estimates for this species. The size of the GRCs in our genome assembly indicates that the two GRCs may have been at least partially assembled separately. We explore this possibility below.

We annotated 41,418 genes in our *B. coprophila* genome assembly: 17,802 on the autosomes, 4,277 on the X chromosome, and 15,812 attributed to the GRCs (**Table 1**). The number of genes that we annotated on the autosomes and X chromosome are comparable to the recently published reference genome for *B. coprophila* [31] (**Table 1**), however, the number of annotated genes overall is greater than in Urban et al. [31]. This is because the reference genome assembly was constructed primarily with somatic tissue sequence (from embryos after GRC elimination from somatic cells), and it is therefore not expected to contain GRC genes.

### GRCs have paralogs throughout genome

To better understand the origins of the GRCs, we conducted reciprocal blast searches with the annotated genes to infer paralogs within our genome assembly. We also conducted a collinearity analysis to identify larger homologous blocks in the genome in which we identified collinear blocks of five or more genes anchored to the reference assembly (for autosomal and X-linked genes) [31] or an assembly we generated with long-read data from male germ tissue (for GRC genes-see **Supplementary Text 2** for methods). This allowed us to increase the continuity of our assembly and to anchor genes within our assembly to known chromosomes (autosomes A-II, A-III, A-IV, and the X chromosome) in the reference genome. From these analyses we wanted to determine 1. Whether the GRC genes have paralogs on other chromosomes in the genome and whether paralogs were mostly on one chromosome, which would allow us to determine the origin of the GRCs, 2. Whether there is evidence for strata on the GRC with different genes having different divergence levels (i.e. some genes older than others) and 3. Whether GRC-GRC reciprocal blast hits are prevalent in the genome assembly, which would give further evidence that the two GRC chromosomes were assembled separately (i.e. that the same gene on homologous GRCs were assembled on separate scaffolds). For convenience, we will call the GRC-GRC reciprocal hits paralogs too, even though the circumstances under which they diverged are not clear.

We found that the GRCs carry many paralogous genes to both autosomes and the X chromosome (**Fig 3A**). Additionally, there is a substantial number of paralogs in which both copies are on the GRC (GRC-GRC paralogs). Overall, 71.4% of the paralogs we identified contained at least one GRC gene. The sequence identity between paralogs showed a unimodal distribution without striking differences between specific paralog groups (**Fig 3B**), suggesting that divergence between paralogs is not dependent on the genomic location of the genes in the paralogs. A collinearity analysis revealed 88 collinear blocks between the GRC and autosomes or the X chromosome, 23 collinear blocks in which both blocks were located on GRC scaffolds, and 5 collinear blocks in which both blocks were located on an autosome or the X chromosome. We anchored 42 blocks to individual chromosomes in the reference assembly and found that the GRCs are homologous to all four chromosomes of *B. coprophila* (**Fig 3C; Supplementary Table 2**), suggesting that the GRCs are not derived from a single chromosome nor from a simple chromosomal rearrangement (e.g. fusion of a chromosomal arm and X chromosome).

**Fig 3.**
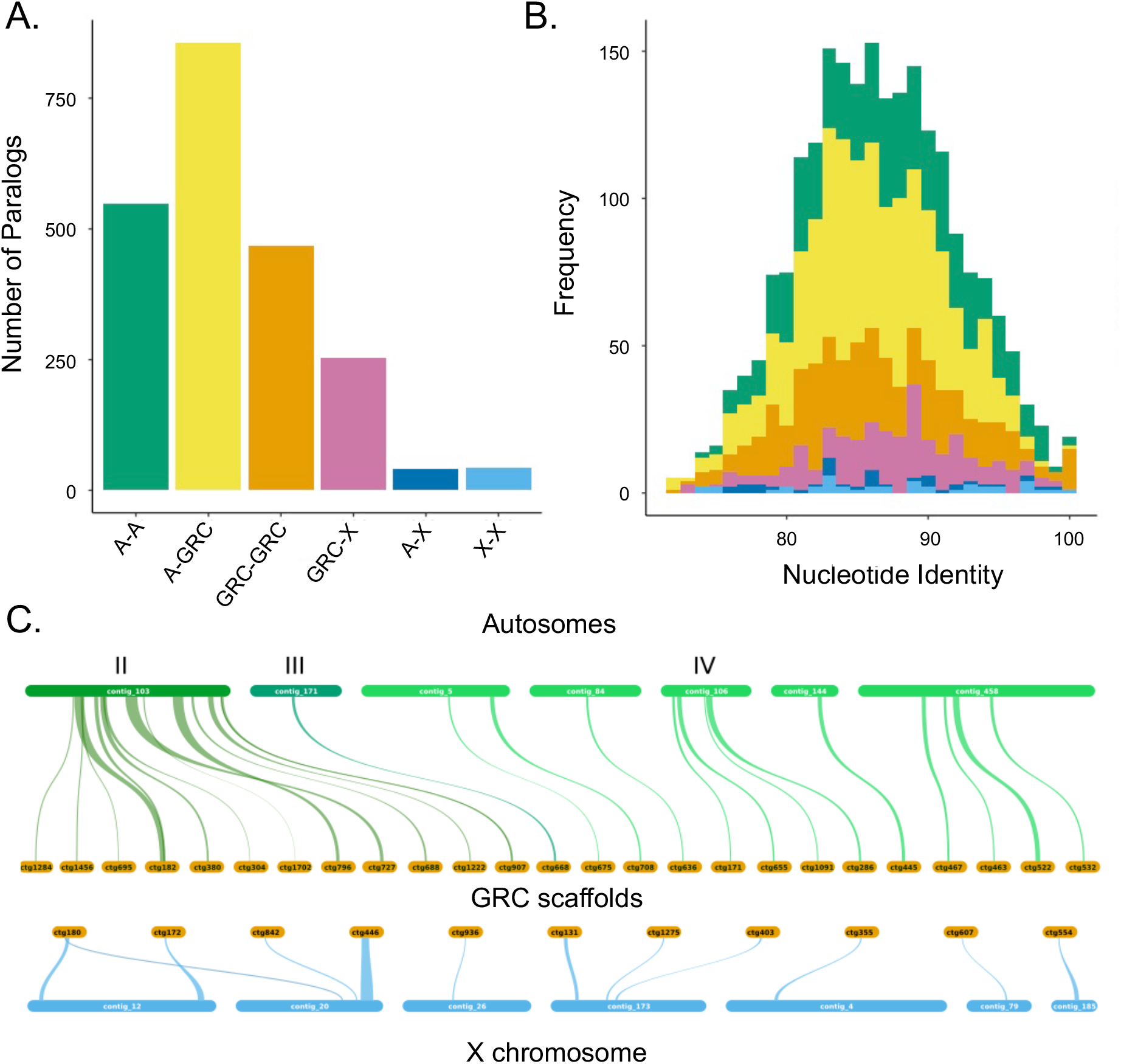
GRC genes have divergent paralogs distributed throughout the core genome. **A.** Number and **B**. nucleotide identity of paralogs between different chromosome types in *B. coprophila*. The majority of paralogs (>70%) involve GRC genes and many paralogs are between the GRC and autosomes. Additionally, all paralog types have similar divergence levels. **C**. Collinear blocks found between GRC scaffolds (orange) and scaffolds anchored to the X chromosome (blue) or individual autosomal chromosomes (A-II, A-III, or A-IV; shades of green). Note that there is variation in the reference assembly in the proportion of scaffolds that are anchored to each chromosome (**Supplementary Table 2**).

The GRC chromosomes in Sciarids were hypothesized to be derived from the X chromosome [23], therefore, one of our aims for the paralogy and collinearity analyses was to test if there is a clear homology between the X chromosome and GRC. Contrary to theoretical expectations, the GRCs carry many paralogous genes to both the autosomes and X chromosome (**Fig 3A**), the divergence between the GRC and X chromosome paralogs was similar to the divergence between the GRC and autosomal paralogs (**Fig 3B)** and we identified collinear blocks between the three autosomes and the GRCs as well as the X chromosome and the GRCs (**Fig 3C**). Therefore, we found no evidence that the GRCs were derived from the X chromosome. Rather, it seems that the GRCs show no clear homology to any specific chromosome, but have homologous regions to all chromosomes in roughly equal proportions. This is similar to recent findings on the GRC in zebra finches, in which it was found that the genes on the GRC in this species also had paralogs located throughout the genome, so there was no clear chromosomal origin for this chromosome. However, in contrast to the zebra finch GRC, where some GRC genes were found to be older than others, the unimodality of divergences of GRC genes to their paralogs in *B. coprophila* suggest the GRC were acquired in a short evolutionary time frame, perhaps during a single event (further explored in the phylogenetic analysis).

In *B. coprophila*, the two GRCs are a homologous pair of approximately 88Mbp (**Table 1**). They form bivalents during female meiosis, but it remains unclear whether the two chromosomes recombine [32]. If the recombination is suppressed, the two GRCs could diverge over time to the extent that the two homologous GRC chromosomes assembled on separate scaffolds. We found that the total size of the GRC scaffolds was about twice as large as we expected given the estimated size of one GRC chromosome (154 Mbp vs. 88Mbp; **Table 1**). This result, in addition to the large number of GRC-GRC paralogs we identified, suggests that the two GRCs indeed are divergent. This suggests that the reciprocal blast hits in which both gene copies were on the GRCs are likely alleles of the same loci on the two homologous GRC chromosomes. However, the GRC-GRC paralogs also show similar divergence distribution as the GRC-autosomal and GRC-X paralogs, suggesting that the two GRCs diverged from each other over extended periods of time (i.e. some genes stopped recombining close to the origin of the GRCs) (**Fig 3B**).

### The two GRCs are heteromorphic and show different sequencing coverage

To further investigate whether the two GRCs are homologous but deeply divergent, we analysed the sequencing coverage of all GRC genes, paralogs in which both copies are on the GRCs, and collinear blocks where both blocks are located on the GRCs. We found the sequencing coverage of GRC genes is bimodal, with two modes at 25x coverage and 30x coverage (**Supplementary Fig 3**). We tested if the two histogram peaks represent genes on the two GRC chromosomes by comparing the coverage of GRC genes in our paralogy analysis in which both genes in the reciprocal blast hit were on the GRC (see above). Indeed, most of these genes have one paralog in the low coverage peak (coverage 18-33x) and the second paralog in the high coverage peak (coverage 23-38x; **Fig 4A**), suggesting that the two GRCs have different sequencing coverages and the GRC-GRC genes in our paralogy analysis are indeed copies of the same gene on different GRC chromosomes. To confirm the association of the two GRC chromosomes with the two coverage peaks we extracted GRC-GRC collinear blocks and their corresponding coverages. Indeed, most of the collinear blocks showed the same pattern -one block containing genes with close to the higher coverage peak and the other genes with a coverage close to the lower coverage peak (**Fig 4B**), however there were a few exceptions to this rule as well (See **Supplementary Fig 4**).

**Fig 4.**
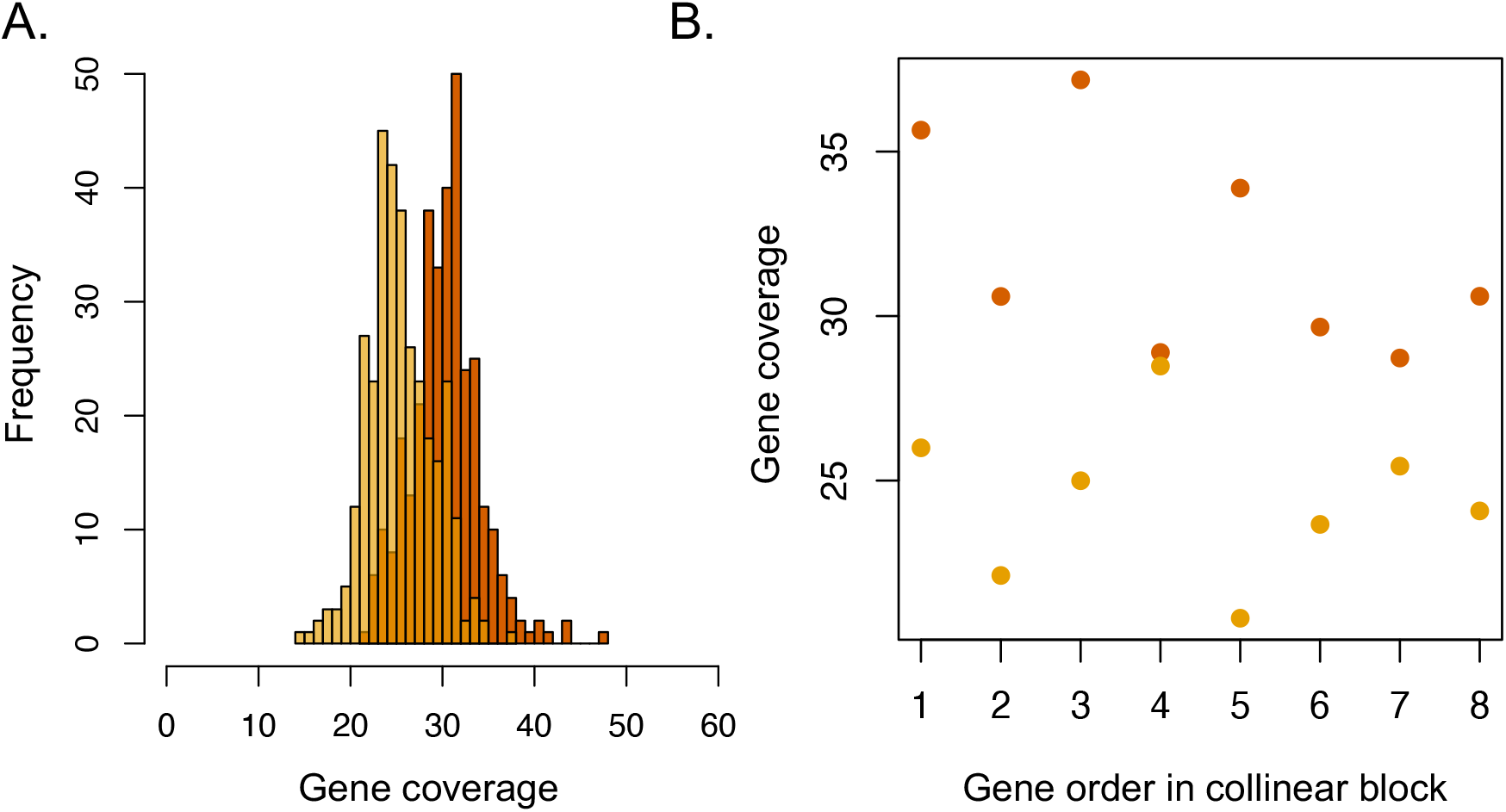
Coverage differences between GRC paralogs. **A**. histogram of coverage differences between GRC-GRC paralogs, the paralog with a higher coverage is included in the darker histogram while the lower coverage paralog is included in the lighter histogram. **B**. One example (out of 23) of a GRC-GRC collinear block comparing coverage of 8 GRC-GRC paralogs. The genes in one collinear block have a higher coverage (∼30-35x coverage) than the other block (∼23-28x coverage).

We were surprised to see that the two GRC chromosomes appear to have different sequencing coverages in male germ tissue. Male germ cells contain two GRCs, and so the heteromorphic GRCs should be at an equal frequency in this tissue. However, males occasionally show variation in the number of GRCs in spermatocytes [24] and our libraries were made from pools of 95 male testes. Therefore, the two GRCs may have been at slightly different frequencies in the flies we sequenced. The differences in GRC frequency in male testes suggests that the variation of GRCs in sperm may not be purely stochastic with respect to the two differentiated GRCs (i.e. one is more likely to be present than the other). However, at present we do not know why one GRC would be more likely to be at a higher frequency than the other. The transmission of GRC chromosomes in *B. coprophila* is unusual: eggs contain one GRC and sperm two, so zygotes initially have three GRC chromosomes (**Fig 1**). Germ cells, however, only contain two GRCs because early in germ cell development one of the three GRCs is eliminated [16]. Until now, it was supposed that this elimination is random, but our data suggests that this cannot be the case, since we would not expect to maintain two divergent GRC homologs if the elimination at this stage was random (i.e. through drift alone). Instead, it seems likely that the elimination of GRCs from early germ cells is likely parent-of-origin specific. Further work is however required to clarify the inheritance of these chromosomes, and whether retention of the two GRCs in early germ cells is non-random with respect to the parent of origin.

### The GRC is old and its evolutionary origins are obscure

In order to better understand how old the GRCs are, we reconstructed the phylogenetic placement of GRC genes in Sciaroidea (the superfamily which contains Sciaridae and Cecidomyiidae, which both carry GRCs, and several other gnat families). We used a set of universal single-copy orthologs (BUSCO) identified in recently published draft genomes for 13 species within Sciaroidea and outgroup species (*Sylvicola fuscatus*) [27] (**Supplementary Fig 5**). We identified 340 BUSCO genes that were duplicated in our *B. coprophila* genome with one copy on the GRC and one copy on either an autosome or the X chromosome (i.e. GRC-A/X paralogs) (**Supplementary Table 2**). We generated a phylogeny from these genes and found that the GRC genes branch within the Cecidomyiidae family; specifically, the GRCs are most closely related to the hessian fly *Mayetiola destructor* (**Fig 5A**). The phylogenetic position of GRC sequences in *B. coprophila* is puzzling, but suggestive of an alternative hypothesis of the origin of GRC to the theory that the GRCs evolved within the Sciaridae family from somatic chromosomes.

**Fig 5.**
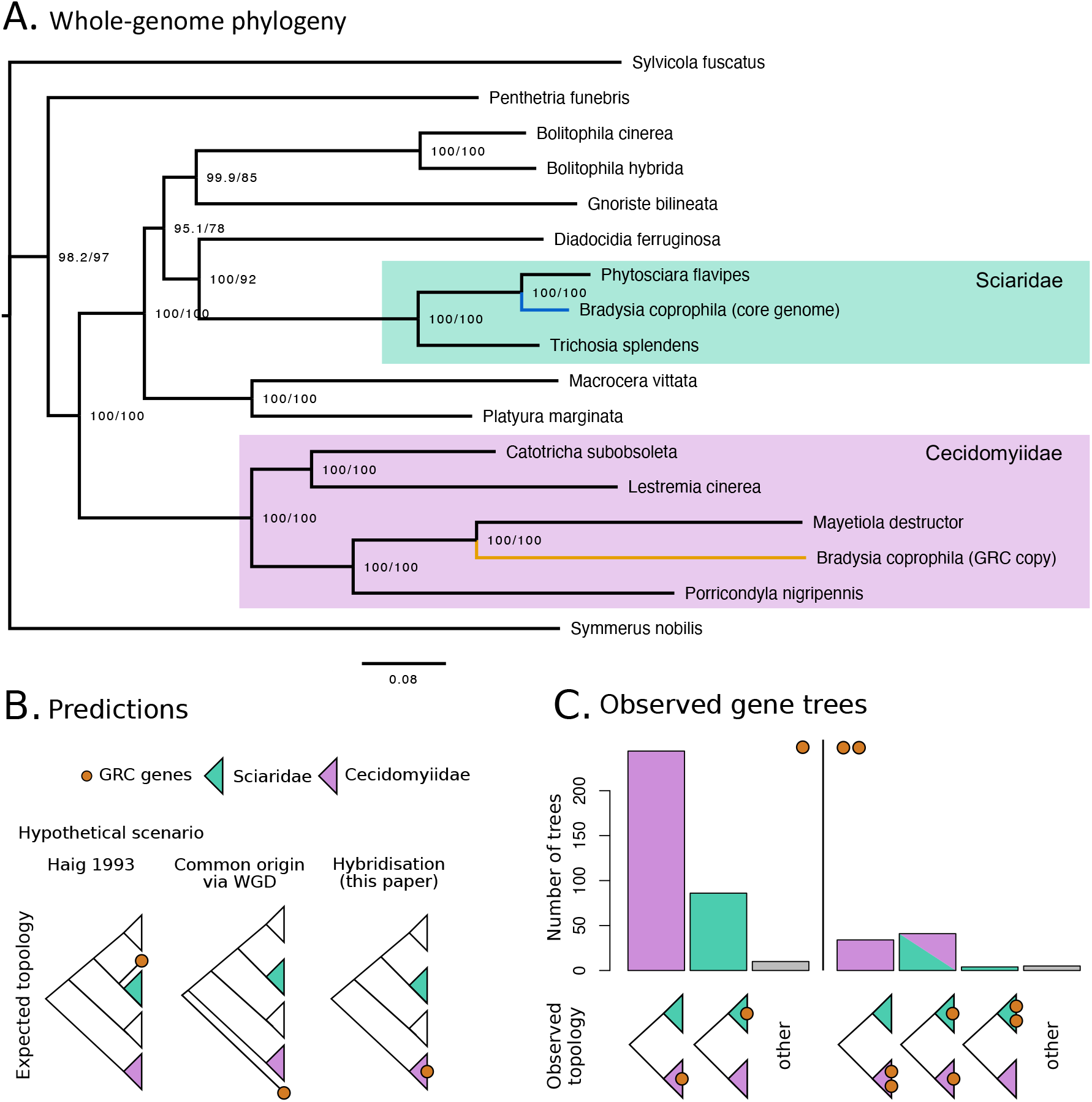
Phylogenetic analysis of conserved genes on GRCs. **A**. Phylogeny generated from 340 duplicated BUSCO genes in *B. coprophila* with one gene copy on the GRC and one copy on either an autosome or the X chromosome. The reconstructed tree identifies the origin of GRC sequences in the Cecidomyiidae family. **B**. Expected gene tree topologies given three hypothetical scenarios: evolution of the GRCs from somatic chromosomes at the root of Sciaridae, common evolutionary origin of GRCs in Sciaridae and Cecidomyiidae through a whole-genome duplication (WGD) event before the split of the lineages, or evolution of GRC in Sciaridae via introgression from Cecidomyiidae. **C**. Breakdown of individual gene tree topologies with respect to position of GRC copies; most of the trees support the hybridization hypothesis (i.e. GRC genes branching from within the Cecidomyiidae). Genes with one GRC gene copy and one copy in the core genome (left side; core gene copy not shown) most commonly have two topologies: GRC copy within Cecidomyiidae (purple) or within Sciaridae (teal), almost no other topologies were found (grey). Genes with two GRC copies and a gene copy in the core genome (right side) frequently have two topologies: both GRC copies within Cecidomyiidae (purple), or one copy within Cecidomyiidae and the other within Sciaridae (striped purple/teal). Four genes also showed a topology with both GRC copies within Sciaridae and only three others showed other topologies (mostly unresolved trees; see **Supplementary Fig 6** for examples of individual topologies).

Instead, our results suggest that the GRCs in Sciaridae originated via introgression from the Cecidomyiidae family, as the GRC branch in the phylogeny falls within the Cecidomyiidae clade, and does not branch from the base of these two clades (which would indicate that the GRCs evolved in the ancestor of Cecidomyiidae and Sciaridae) (**Fig 5B**). This raises questions about how these chromosomes evolved. The most parsimonious explanation from our phylogenetic data is that the GRCs in Sciaridae arose through a hybridisation event between early Sciarids and Cecidomyiids, as the *B. coprophila* GRC branch falls within Cecidomyiidae, but is longer than the root of Sciaridae family, suggesting the hybridisation event has probably happened prior to diversification of the Sciaridae family. To explore the hypothesis of GRC origin through hybridisation in Sciaridae, we examined all gene trees in which one *B. coprophila* gene was located either on an autosome or the X chromosome and one or two genes were located on the GRC. We found that most of the gene trees support the hybrid origin hypothesis (**Fig 5C**). In 410 of 424 (97%) gene trees, the autosomal/ X linked gene copy fell within the Sciaridae clade, as expected. For single copy GRC paralogs, 71.8% (244), were identified as members of Cecidomyiidae family and in a minority of these trees the GRC gene fell within the Sciaridae (25.3%; 86) (**Fig 5B**). The terminal branches of GRC genes within the Sciaridae family are significantly shorter compared to those within the Cecidomyiidae family (mann-whitney p-val < 0.0001; **Supplementary Fig 7**). Hence we hypothesise the GRC genes within Sciaridae likely represent more recent acquisitions on the GRCs from core chromosomes within the Sciaridae, which is not unexpected as the GRC genes have likely been present in Sciaridae for more than 44 million years [33–35]. For BUSCO gene trees in which two gene copies were on the GRC and one was on an autosome or the X chromosome (84 genes), 41.7% (35) had a topology where both GRC genes fell within the Cecidomyiidae, 50% (42) had a topology where one gene fell within the Cecidomyiidae and one fell within the Sciaridae, and a much smaller proportion had both genes branching from within the Sciaridae (4) (**Fig 5B**). Overall, these results strongly support the hypothesis that the GRCs within Sciaridae arose through introgression from the Cecidomyiidae, perhaps through a hybridization event somewhere near the base of the Sciaridae.

The results of this study raise many questions about the evolution of GRCs in Sciaroidea (both Cecidomyiidae and Sciaridae). Our study rejects the hypothesis that GRCs in Sciaridae arose from the X chromosome in this lineage, and instead suggests that they arose through introgression from Cecidomyiidae, perhaps through an ancient hybridisation event. There are very few examples where interspecies crosses gave rise to additional chromosomes with non-Mendelian inheritance, with one exception being the PSR (paternal sex ratio) chromosome in the parasitic wasp *Nasonia* [36,37]. The PSR chromosome is a B chromosome that interferes with sex determination in its wasp host and is thought to have evolved through hybridization with a parasitoid wasp in the genus *Trichomalopsis* [37]. GRCs are present in both Cecidomyiidae and Sciaridae, but are thought to have evolved independently and are not thought to be present in other Sciaroidea families [18,38,39]. It is tempting to speculate that the GRCs in Sciaridae and Cecidomyiidae share a common origin, however, we currently do not have GRC sequence from species within Cecidomyiidae to assess this idea. Such a dataset would be extremely useful to establish whether the GRCs in *B. coprophila* show greater homology to the Cecidomyiid GRC genes, or their autosomal counterparts as this analysis only took into account somatic gene sequence in all Cecidomyiid species.

In many ways, GRCs in Cecidomyiidae are quite different from those in Sciaridae: they are much more numerous, are generally exclusively maternally transmitted, and are smaller than those in Sciaridae [18]. Since the GRCs in Cecidomyiidae are numerous, they were originally thought to have evolved through multiple rounds of whole genome duplication, followed by restriction of the duplicated chromosomes to the germline (although note that this idea is somewhat controversial as the GRCs have different banding patterns to the core chromosomes) [18,40]. If the GRCs in Sciaridae arose through hybridisation with Cecidomyiidae, GRCs in both lineages would have evolved through polyploidisation, although via quite different routes and with different evolutionary trajectories after the establishment of GRCs. It is a striking coincidence that the presence of GRCs in Sciaroidea is associated with unconventional non-Mendelian reproduction systems in both the Cecidomyiidae and Sciaridae. Future studies will establish whether this is truly a coincidence, whether the unconventional transmission dynamics in both families somehow facilitates the evolution of GRCs or vice versa. For instance, the fact that the GRCs in Sciaridae are eliminated from somatic cells in much the same way as the X chromosome is eliminated for sex determination is suggestive that either the GRCs have become established in the germline by manipulating the mechanism of sex determination, or that the system of sex determination in Sciaridae arose through manipulating the mechanism by which GRCs are eliminated from somatic cells. However, we need to learn much more about the genetic underpinnings of sex determination in these clades, and to establish the timing of the evolution of different parts of the chromosome system in these families to establish how/whether GRC evolution and the evolution of the unusual sex determination mechanism in Sciaridae (and Cecidomyiidae) are related.

### Function of GRCs in Sciaridae

There has historically been some debate as to whether the GRCs in Sciaridae provide any sort of necessary function [41]. The GRCs in *B. coprophila* are primarily heterochromatic, as evidenced by cytological studies showing that they are densely staining over much of *B. coprophila* development, and possess modifications that are characteristic of constitutive heterochromatin [24,42]. It has been hypothesized that *B. coprophila* GRCs might be transcribed in the germline at 96 hours after oviposition, when they become euchromatic [24] and perhaps also during interphase between male meiosis I and II or after male meiosis in a related Sciarid, *Trichosia* [43]. Since heterochromatin is gene-poor, it was thought that few if any genes reside on the GRCs, similar to many B chromosomes, which often contain an excess of satellite DNA [36,44]. However, to the contrary, the sequence data presented here have revealed that there are many genes on the *B. coprophila* GRCs and they are paralogs of genes on the other chromosomes. Recently it has also been reported for other plants and animals that genes on eliminated DNA have paralogs in the other chromosomes [10,45]. Although it remains to be seen whether the multitude of *B. coprophila* GRC genes are transcribed and play an important role, with GRC genes now identified, future studies can elucidate when and where their transcription occurs and determine whether these chromosomes are necessary in *B. coprophila*.

Some evidence has suggested that Sciarid GRCs may play a role in reproduction, specifically in sex determination. *Bradysia coprophila* and many other species of Sciarid flies are monogenic, where mothers have only sons or only daughters. This trait is only found in some Sciarids and seems to be correlated with the presence of GRCs. Indeed, all Sciarid species that are monogenic have GRCs, suggesting that these GRCs might play a role in sex determination. Additionally, a strain of *Bradysia impatiens*, which is a monogenic species with GRCs, arose in the laboratory which became digenic, and this was correlated with the loss of GRCs [32]. Therefore, GRCs may be similar to the PSR B chromosome in the jewel wasp *Nasonia vitripennis* which causes female-to-male conversion; a transcript from a gene on the PSR chromosome has been identified which causes this effect [46]. However, the link between GRCs and sex determination is not air-tight, as Sciarid species that are digenic (i.e. females produce offspring of both sexes) can either have GRCs or lack these chromosomes [32]. It is of course possible that the gene(s) for the monogenic trait have been lost from GRCs in the digenic Sciarid species that retain GRCs. More research on this topic is needed to establish whether GRCs do have a function relating to sex determination in Sciaridae.

### Concluding remarks

*Bradysia coprophila* has a fascinating chromosome inheritance system, which displays several examples of non-Mendelian transmission and contains two germline restricted chromosomes. Understanding more about how this system evolved can tell us about the evolution of alternative non-Mendelian reproduction systems as well as about the evolution of germline restricted chromosomes and germline soma differentiation. Through sequencing the germline restricted chromosomes in the Sciarid *B. coprophila*, we have determined that the two germline restricted chromosomes in this species contain many protein coding genes. Additionally, the two GRCs in *B. coprophila* seem to form a non-recombining chromosome pair, with divergent homologs on the two GRCs. Although much still needs to be elucidated about how these chromosomes are transmitted, this is one of the only examples of heteromorphic chromosomes which are not sex chromosomes. For this reason, these chromosomes provide food for thought, as we can explore whether their evolutionary trajectory has followed that of heteromorphic sex chromosomes.

Additionally, our results indicate that the origin of the GRCs in *B. coprophila* is through introgression from Cecidomyiidae, a gall gnat family also in the infraorder Bibionomorphia which also displays a non-Mendelian inheritance system and GRCs. This is a fascinating example of cross-family introgression. Using a time calibrated phylogenetic tree, we roughly estimated that the hybridisation happened 116 -50 mya, and between 31 -97 my after split of the two ancestors of Sciaridae and Cecidomyiidae (See **Supplementary Text 3** for details). Although animals of similar divergence have been successfully hybridised in the lab [47], we present the first evidence for a cross-family hybridisation event in nature with evolutionary consequences. Gene flow between very divergent lineages seems to be frequently associated with polyploidisation (for example in burrowing frogs [48], or Arabidopsis [49]), supporting our view that GRCs evolved in the current form a whole genome introgressed from the ancestor of Cecidomyiidae.

Finally, our results add additional insight into the evolution of germline restricted DNA. Studies on germline restricted DNA in taxa with chromatin diminution (i.e. portions of chromosomes rather than whole chromosomes are restricted to the germline) suggest that this system evolves to resolve germ/ soma conflict over gene expression. However, our results strongly suggest that the GRCs in *B. coprophila* evolved not as a means to resolve germ/soma conflict, but likely instead to resolve conflict between chromosomes which were introgressed into Sciaridae from Cecidomyiidae. The GRCs in zebra finches, as well, are not suggested to have evolved as a means to resolve germ/ soma conflict, but are instead proposed to have evolved from a selfish B chromosome [13]. Investigating the evolution of GRCs in more lineages will help to settle this question, but it seems that the origin of GRCs are likely to be different than the origins of germline restricted DNA in systems with chromatin diminution, and it may be useful to consider the evolutionary pressures which lead to these two systems separately. However, after germline restricted DNA evolves, it might follow a similar evolutionary trajectory in both chromatin diminution and chromosome elimination systems, given that in both systems researchers have found that germline restricted DNA are enriched for genes that function in germline maturation/ function [8–10]. Understanding more about whether GRC genes are expressed in *B. coprophila*, and how/ whether they have a germline related function, will provide additional insight into how different types of germline restricted DNA are related, and whether GRCs in *B. coprophila* provide a similar function to other lineages with GRCs.

## Materials and Methods

### Fly culture maintenance

*Bradysia coprophila* lines used in this study have been maintained in the laboratory since the 1920s [28]. Most of the biological literature refers to this fly as *Sciara coprophila*, although the genus name was changed from *Sciara* to *Bradysia* some decades ago [50]. We refer to it here as *Bradysia coprophila*, but *Sciara tilicola* (Loew, 1850), *Sciara amoena* (Winnertz, 1867) and *Sciara coprophila* (Lintner, 1895) are all synonyms. Our *B. coprophila* cultures were obtained from the Sciara stock centre at Brown University and kept at the University of Edinburgh since October 2017. We maintain colonies by transferring one female and two males to a glass vial (25mm diameter x 95mm) with bacteriological agar and allowing the offspring of the female to develop. During development, we add a mixture of mushroom powder, spinach powder, wheat straw powder and yeast to the vials two to three times a week until the larvae pupate.

### gDNA extractions and sequencing

We sequenced genomic DNA from somatic (heads) and germ (testes and sperm) tissue of 1-2 day old adult males. We generated Illumina short read data from somatic and germ tissue. We dissected males which had been put on ice in a vial (to slow down males) on a clean slide in a dish of ice under a dissecting scope. For the dissections, we used jewellers forceps to separate the head from the body and then placed the head in a 1.5ml microcentrifuge vial on dry ice. We then placed a drop of sterile 1X PBS on the body of the male and used forceps and insect pins to slowly pull the claspers away from the body until the claspers and male reproductive tissue separated from the body. We then severed the ejaculatory duct and placed the testes in a separate microcentrifuge tube. We collected males over several days and stored the samples at -80°C until DNA extractions, sequencing a pooled sample from the tissue from 95 males.

The DNA extraction protocol we used was a modified version of the Qiagen DNeasy Blood and tissue kit extraction procedure (see **Supplementary text 1** for full protocol). We quantified DNA on a qubit fluorometer (v3). We sequenced the samples on the Illumina Novaseq S1 platform, generating PE data with 150bp reads and 350bp inserts through Edinburgh Genomics.

### Genome assembly and annotation

We generated a genome assembly with both the somatic and germ tissue short read libraries (**Supplementary Table 1**). We also generated a genome assembly from long read sequence data from germ tissue, but the short-read assembly produced a more complete genome assembly according to BUSCO gene assessments, so this assembly was used for gene annotation. We used the long read assembly for the collinearity analysis to increase the continuity of GRC scaffolds (See **Supplementary Text 1** for details).

For the short read libraries, we trimmed the raw reads with fastp with parameters --cut_by_quality5 --cut_by_quality3 --cut_window_size 4 --cut_mean_quality 20 [51], and used fastqc to investigate read quality (https://www.bioinformatics.babraham.ac.uk/projects/fastqc/). We generated an initial assembly with CLC assembly cell using default settings (Qiagen-v 5.0.0), then used blobtools [52,53] to investigate contamination in the raw reads (See **Supplementary Fig 1** for blobplot), using bamfilter to retain reads which had a GC content between 0.14 and 0.51 and a coverage higher than 7 (which excluded most Prokaryotic sequences identified as contaminants). We generated an assembly with spades [54] using the filtered reads and k-mer sizes of 21, 33, 55, and 77. We conducted a BUSCO analysis (version 4.0.2) [55] using the insecta database (insecta_odb10) to assess whether single copy orthologs expected to be present in insect genomes are present in our draft genome. We then annotated the genome using the braker2 pipeline [56], aligning RNAseq reads from male and female germ tissue to the genome using Hisat2 (using default settings, v2.1.0) [57], and using RepeatModeler (v2.0.1 using default settings) [58] and RepeatMasker (v4.1.0) [59] with the RepeatModeler output and known insect repeats as the repeat library, and the settings -gff -gc 35 -xsmall -pa 32 -no_is -div 30 to mask the genome assembly.

### Identification of GRC scaffolds

We used a combination of two techniques to identify scaffolds belonging to the GRC in our assembly. One technique employs coverage differences between the germ and somatic tissues to identify which chromosome a scaffold belongs to. Since the number and type of chromosomes differs between the somatic and germ tissues (**Fig 2A**), we expect autosomal scaffolds to have a log2 coverage difference (germ/soma) of approximately -1 (i.e. at 2X the frequency in somatic tissue compared to germ), X-linked scaffolds to have a coverage difference of approximately 1, and GRC scaffolds to have very few reads mapping to them from the somatic library but a diploid coverage level in the germ tissue library. We mapped the germ and somatic reads to the genome assembly with bwa mem (v0.7.17) using default settings and counted the number of reads from each library mapping to each scaffold [60]. Due to somatic contamination in the germ library, the coverage differences displayed the pattern we expected and we were able to distinguish autosomal and X linked scaffolds but the autosomal and X chromosome scaffolds had slightly different coverage differences than expected. We labelled scaffolds with a coverage difference of >-1 to <-0.1 as autosomal, those with a coverage difference of >-0.1 to <0.5 as X-linked, and scaffolds with a coverage difference greater than 0.5 as GRC linked (**Fig 2B)**.

The second technique we used to assign scaffolds to chromosomes utilizes differences in the frequency of k-mers in the raw sequencing reads of each library. We used the kat comp command (kat v 2.4.1) [61] to generate a 2D histogram comparing 27-mer composition between the germ and somatic libraries (**Fig 2C**). We extracted 27-mers and their coverages using kmc dump [62]. Using custom scripts we assigned k-mers with a frequency between 125 and 175 in the somatic library and between 80 and 140 in the germ library as autosomal, k-mers with a frequency between 50 and 100 in the somatic library and 60 and 100 in the germ library as X-linked, and k-mers with a frequency <5 in the somatic library and >10 in the germ library as belonging to the GRC. We searched for exact matches these k-mers in the assembled scaffolds using bwa mem (v0.7.17 with -k 27 -T 27 -a -c 5000 parameters) [60] and generated a score comparing the number of k-mers mapping to each scaffold from the chromosome type with the most k-mers mapping to the scaffold by the length of the scaffold (see **Supplementary Fig 2** for plots assessing the efficiency of the k-mer identification technique). Scaffolds with mostly autosomal k-mers mapping to them and a score greater than 0.4 were assigned as autosomal. Similarly X-linked scaffolds with a score greater than 0.4 were assigned as X-linked, and GRC scaffolds with a score greater than 0.8 were assigned as belonging to the GRC chromosomes (**Supplementary Fig 2**). We then compared the scaffolds assigned using the k-mer and coverage techniques. Only scaffolds that were assigned as the same chromosome type with both techniques were included in downstream analyses.

### Genome wide paralog identification

We conducted an all-by-all blast search of annotated genes to identify gene paralogs in our assembly both using nucleotide sequences (**Fig 3A/B**) and translated amino acid sequences (**Fig 3C**). First, we extracted transcripts for each gene with gffread (v0.11.7) [63], and used the longest transcript for each gene as the gene sequence. We identified paralogs using reciprocal blast of translated genes with an e-value cutoff 1e^-10 and reciprocal hits that span at least 70% of both genes. Then, for the collinearity analysis, we mapped GRC-linked genes to the long read assembly (**Supplementary Text 1**), and autosomal and X linked genes to the reference assembly (NCBI accession: GCA_014529535.1 [31]) using blastn with an e-value cutoff of 1e^-10 (2.5.0+). Using the mapped set of genes and the amino acid reciprocal blast, we performed a collinearity analysis using MCScanX with default parameters (at least 5 colinear genes, genes must match the strand). Note that in the reference assembly 20-46% of A-II, 8-19% of A-III, 37-52% of A-IV, and 93-100% of the X chromosomes are anchored (**Supplementary Table 2**) [31]. The synteny blocks between GRC scaffolds and individual anchored autosomal and X scaffolds respectively were visualized on **Fig 3C** using SynVisio (commit 4a4361f, [64]).

### Coverage analysis of paralogs

We used BEDtools coverage with settings -mean -a to compute the mean coverage across each annotated gene within the *B. coprophila* genome assembly (v2.26.0) [65]. We examined the histogram of mean coverages of all GRC linked genes, then examined the subset of GRC genes which were identified in the paralogy analysis as being involved in GRC-GRC paralogs. In order to explore whether these genes are alleles of the same gene on different GRC homologs or true GRC-GRC paralogs, we identified which gene in each GRC-GRC pair had a higher coverage, and plotted it in a separate histogram from the lower coverage gene in the same pair.

We also examined the coverage of the GRC-GRC collinear blocks identified through the collinearity analysis. We identified 23 GRC-GRC collinear blocks and compared the coverage of each paralog along the block. We only took into account genes with a mean coverage less than 45x, as the majority of GRC linked genes had a coverage between 15x and 45x (see **Supplementary Fig 3**). We identified how well each of the collinear blocks met the expected coverage patterns (i.e. one block having a higher coverage than the other) by computing a statistic comparing the number of genes that meet the expected coverage patterns by the total number of genes in the collinear block (**Supplementary Fig 4**).

### Phylogenetic analysis of the GRCs origin

We utilized draft genome assemblies for 14 Sciaroidea species and 2 species outside the Sciaroidea, most of which we obtained from Anderson et al. [27] with the exception of *Mayetiola destructor*, which we obtained from NCBI (accession: GCA_000149195.1). We conducted a BUSCO analysis (version 4.0.2) [55] using the insecta database (insecta_odb10) on each genome assembly, along with our *B. coprophila* assembly, to identify single copy orthologs in each genome. We excluded the *Exechia fusca* genome from further analyses as this genome had a low proportion of complete BUSCO genes identified, indicating that the genome was likely of poor quality. We identified the chromosomal locations of each BUSCO gene identified in the *B. coprophila* assembly and extracted the BUSCO IDs for all genes which were duplicated and had one gene copy on an autosome or the X chromosome and either one or two gene copy on the GRCs (**Supplementary Table 3**). We took the amino acid sequence of these BUSCO genes for *B. coprophila* (all copies) and the longest amino acid sequence for each BUSCO ID per species as the gene sequence in the genome assemblies from all other species (although note that most of the other Sciaroidea species had relatively low rates of gene duplication--See **Supplementary Fig 5**). We only retained BUSCO IDs in the analysis in which 80% of the species of interest had complete versions of the gene.

With the 340 remaining BUSCO IDs with one somatic gene copy in *B. coprophila* and one GRC gene copy, we reconstructed a phylogeny in IQtree using settings -alrt 1000 –bb 1000 (v2.0.3) [66–68]. We also calculated gene trees for each individual BUSCO ID for the 340 IDs mentioned above as well as for 84 BUSCO IDs which had one somatic gene copy in *B. coprophila* and two GRC gene copies using the same settings. We wanted to determine how many individual gene trees support the position of the GRC branch in the concatenated phylogeny, so we used a custom script to summarize for each gene whether the GRC gene copy was found in the Cecidomyiidae clade, the Sciaridae clade, or at some other location in the phylogeny.

## Acknowledgements

We would like to thank members of the Ross lab for comments on this paper. We would also like to thank Natália Martínková, Stuart Baird, Alex Suh and the rest of the GRC community for providing feedback on this work. Thanks to John Urban for providing access to the *B. coprophila* reference genome ahead of publication. CH would like to thank NSERC and the Darwin Trust of Edinburgh for postgraduate financial support. LR would like to acknowledge funding from the European Research Council Starting Grant (PGErepo) and from the Dorothy Hodgkin Fellowship DHF\R1\180120. Financial support from NIH/GM121455 to SAG is gratefully acknowledged.

## Data Availability

Sequence read data will be submitted to NCBI under accession number XXXX. The repository https://github.com/RossLab/Bradysia-GRCs contains scripts associated with this project.

