## Supplemental for "Evolution of gene-rich germline restricted chromosomes in black-winged fungus gnats through introgression (Diptera: Sciaridae)"

**Supplementary Text 1: Detailed description of the chromosome inheritance system in *Bradysia coprophila*.**

The chromosome system in *B. coprophila*, and in sciarids generally, is unique in several ways including chromosome transmission patterns, sex determination, and the presence of GRCs (see **Fig 1** for transmission patterns). All sciarids studied to date have a system of reproduction known as paternal genome elimination, where males only transmit maternally inherited chromosomes to offspring [1,2]. Paternal genome elimination has evolved independently in at least seven arthropod lineages, including the related gall gnat family Cecidomyiidae [3]. In all species with paternal genome elimination, meiosis occurs in a Mendelian manner in females, but in males meiosis is aberrant. In male meiosis in sciarids, there is a monopolar spindle in meiosis I. Maternally inherited chromosomes move towards the monopolar spindle, while paternally derived chromosomes move away from it and are discarded in a bud of cytoplasm [2]. Thus, only the maternal complement of chromosomes is transmitted to the sperm. This phenomenon in *B. coprophila* was the first example of “imprinting”, to our knowledge, by which the cell recognizes the maternal or paternal origin of a chromosome [4]. Interestingly, the GRCs always segregate with the maternal set of chromosomes. Therefore, all of the GRCs (typically two in *B. coprophila*) are transmitted through sperm, regardless of whether they are of maternal or paternal origin [4]. This is one of the few examples of chromosomes which seem to evade paternal genome elimination. In the second division of meiosis in *B. coprophila* there is a bipolar spindle, however there is a nondisjunction of the maternal X chromosome in this division such that only one sperm develops through male meiosis. This sperm contains a haploid set of autosomes, typically two GRCs, and two X chromosomes [1,2]. There is some variation in the number of GRCs in each sperm, ranging from 0-4 in *B. coprophila* [4]. Variation in GRC number is thought to be due to nondisjunction events which can occur in early germ cell divisions, however, the majority of sperm (78%) carry two GRCs [4]. In female meiosis, the GRCs form a bivalent during meiosis, and one GRC segregates into each egg (i.e. meiosis is typical) [4].

As a result of the unusual type of meiosis in male sciarids, *B. coprophila* zygotes typically carry a diploid set of autosomes, three X chromosomes (one inherited from their mother and two from their father), and three GRCs. All sciarids have XO sex chromosome system (i.e. males are XO and females are XX, and there is no Y chromosome), but sex is determined via X chromosome elimination from somatic cells early in development. In the 7-9 cleavage division, either one X chromosome (for females) or two X chromosomes (for males) are eliminated from somatic cells [5]. Elimination occurs due to a failure of separation of the sister chromatid arms during mitosis, resulting in the chromosomes being left on the metaphase plate and not being incorporated into daughter nuclei [6]. It is thought that the number of X chromosomes eliminated is maternally controlled, since *B. coprophila* females are monogenic, and produce exclusively female or male progeny [7]. Females that produce female offspring carry a large inversion on the X chromosome that is always associated with female-producing females [2,8]. GRC elimination from somatic cells occurs in a remarkably similar manner, with the exception that GRC elimination occurs in the 5-6 cleavage division and all GRCs are eliminated from somatic cells [5].

In germ cells, there is also an elimination of one X chromosome and typically one GRC. In this case, elimination occurs in a somewhat mysterious manner in early germ cell development, when one X chromosomes and all but two GRCs are eliminated by being ejected from the germ cell through a cytoplasmic bud [9,10]. Therefore, early germ cells of both males and females in *B. coprophila* have the same chromosome constitution, with a diploid set of autosomes, X chromosomes, and GRCs. This mechanism also regulates the number of GRCs and prevents their accumulation over time, as all but two GRCs are always eliminated from early germ cells.

Less is known about the mechanism of the chromosome system in other sciarid species, but across the family all species studied exhibit paternal genome elimination and X chromosome elimination early in development as the means of sex determination. Although only a handful of species have been studied in detail, evidence suggests that most, but not all sciard species carry GRCs, with the number of GRCs ranging from 0-4 [2]. The two species in which GRCs are absent are closely related to each other, suggesting that GRCs were likely lost in these species. Additionally, monogeny, or females that produce offspring of only one sex, is present in some, but not all species across Sciaridae [2]. There seem to be many transitions in this trait across Sciaridae, with some species being monogenic, some being digenic (i.e. females produce offspring of both sexes), and some species having a mix of these two types of females. Very little is known about the genetic underpinnings of this trait.

Overall, the evidence suggests that paternal genome elimination and X chromosome elimination as a means of sex determination evolved once at the base of Sciaridae. It is less clear how GRCs and monogeny evolved. It was originally suggested that the presence of GRCs and monogeny are related, as *Bradysia ocellaris,* a species that has lost GRCs is digenic. Additionally, a lab line of *Bradysia impatiens* that was bred to lose GRCs transitioned from monogenic to digenic reproduction [4]. However, these facts are anecdotal and there are also several species with digenic reproduction that do carry GRCs (reviewed in [2]).

Cecidomyiidae, gall gnats also in the Infraorder Bibionomorpha, have a similar reproduction system to Sciaridae, in that both families exhibit paternal genome elimination and X chromosome elimination as a means of sex determination, GRCs, and a mix of monogenic and digenic species [11]. However, cecidomyiid species have two X chromosomes (i.e. females are $X_1X_1X_2X_2$ and males are $X_1X_2OO$), and the factor that controls X chromosome elimination in offspring is associated with an inversion on an autosome (rather than the X chromosome in *B. coprophila*) [12]. Additionally, GRC characteristics are quite different in this family, with species containing many small GRCs, which are maternally transmitted and do not seem to form bivalents during female meiosis.

### Supplementary Text 2- Supplementary Methods

*DNA extraction procedure*

For gDNA extractions, for both short read and long read libraries we followed a similar protocol. All the centrifugation steps took place at 4°C and 13,000rpm, unless otherwise stated. Tissue samples were stored at -80°C until DNA extractions. Before extraction, we briefly froze the samples in liquid nitrogen and crushed the tissue with a micro-pestle. We then added 360µl of Cell Lysis Buffer (Qiagen) with 40µl of Proteinase K (20 mg/ml) (Qiagen), and incubated overnight in a shaking incubator at 55°C. We then added 4µl of RNase A (100 mg/ml), mixed by inverting the sample tube, and incubated the sample for 1 hour at 37°C. We cooled the sample on ice for 5 minutes, then added 133 µl of Protein Precipitate Buffer (Qiagen), mixed by gently vortexing the sample and incubated on ice for 10 min. We then centrifuged for 15 min at 4°C, transferred the supernatant to a new tube containing 400µl isopropanol, and mixed by inversion. For the short read samples, we then incubated the sample overnight at -20°C, while for the long read samples, we incubated the sample for 10 min at room temperature. We then centrifuged the sample for 20 min, and discarded the supernatant by inverting the tube. We washed the DNA pellet twice with 300µl freshly prepared 70% EtOH, then centrifuged the sample for 20 min, and carefully removed the supernatant by pipetting. We air dried the DNA pellet for approximately 30 min, and resuspended the pellet in 60µl TE after it dried.

*Long-read assembly*

In addition to the short read sequencing data, we also extracted DNA (using the protocol above) from approximately 250 male testes to generate long read germline data. We sequenced the sample at Liverpool Genomics using a low input library prep procedure and PacBio sequencing on 3 SMRT cells. We used red bean (previously known as wtdbg2) with the parameters -L 1000 -x sq for the initial genome assembly (v2.5) [13], then polished the assembly three times with the long read library using minimap2 with parameters -c -x map-pb to map the long reads to the assembly (v2.17-r941) [14] and racon with parameter -u to polish the assembly (v1.4.10) [15]. We then polished the assembly twice with short read data (with only germline the library) using minimap2 and polishing with racon (see **Supplementary Table 1** for assembly statistics).

The long read assembly, compared to the short read assembly, showed two unfortunate problems: much lower mapping rates of GRC k-mers (**Supplementary Fig 2B**), and lower BUSCO score (BUSCO score: 93.6% complete BUSCOs vs. 98.3% for the short read assembly). We suspect the problems stemmed from high error profiles of long reads combined with high levels of paralogy across the genome hindering precise genome polishing and subsequently leading to frame shifts in gene models. This problem could be resolved in the future with newer sequencing approaches, such as HiFi reads with much smaller error rates, or Haplotagging. However, the long read genome assembly still featured much higher continuity (N50: 576,242 vs 18,920 for short-read assembly). Therefore, we used the short read assembly for annotation and gene level comparisons but used the long read assembly to link individual GRC genes found in the short read assembly for the collinearity analysis (**Fig 3C**).

### Supplementary Figures/ Tables

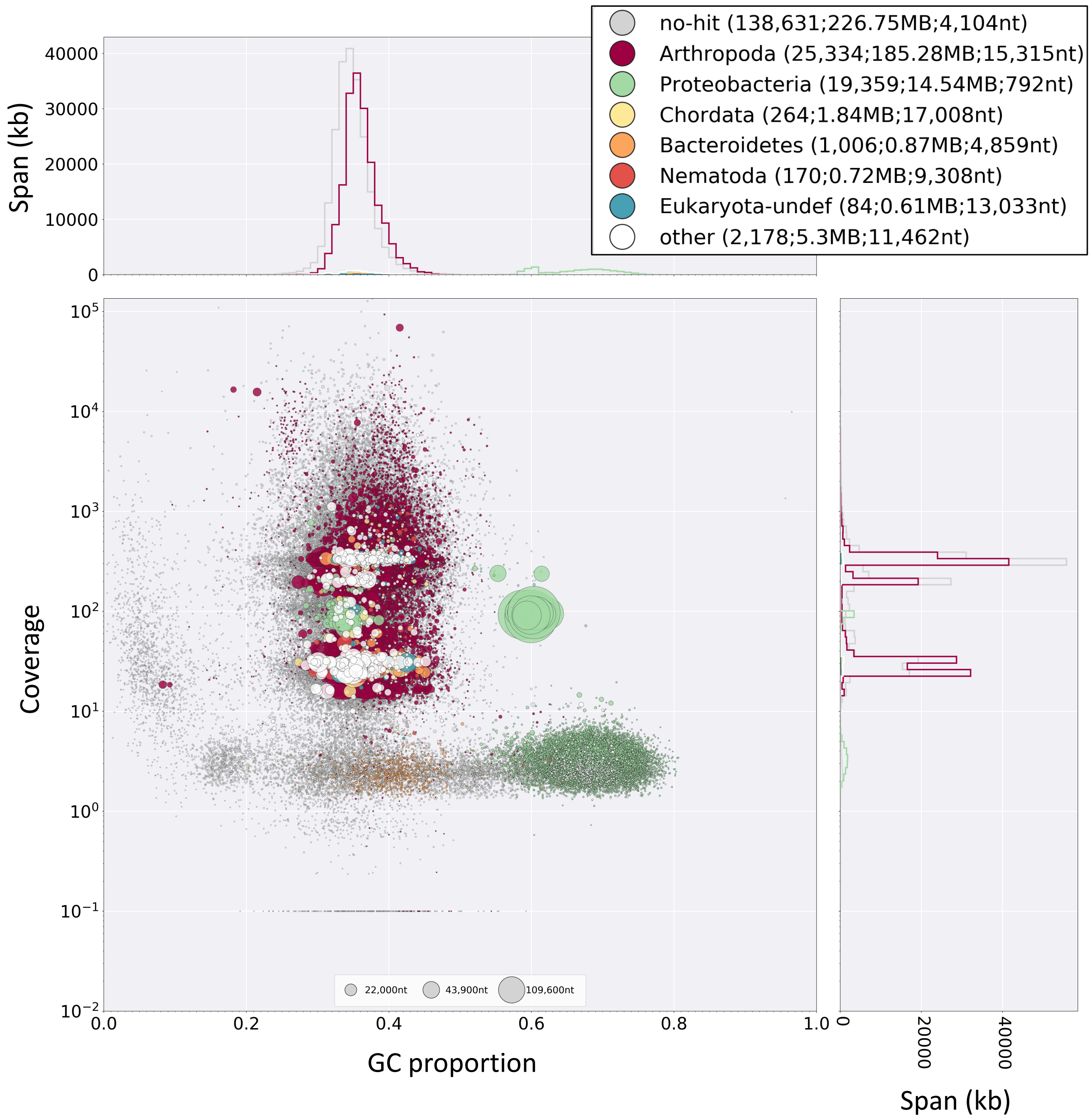


**Supplementary Fig 1. Blobplot of unfiltered assembly** generated from both germ and somatic libraries showing scaffold coverage vs. scaffold GC (size of dot indicates scaffold size and colour taxonomic assignment). Reads mapping to scaffolds with a GC content between 0.14 and 0.51 and a coverage higher than 7 were retained for the final assembly.

**Supplementary Table 1. Summary statistics for the short read and long read assemblies used in this study.** The short read assembly was used for gene prediction as it was closer to the expected genome size compared to the long read assembly and also was more complete according to BUSCO assessment.

| | Short read (Illumina) | Long read (PacBio) |
|---|---|---|
| Size | 398 MB | 415 Mb |
| # scaffolds | 46,532 | 3505 |
| N50 | 18.9 Kb | 576 Kb |
| L50 | 5203 | 135 |
| GC | 35.4% | 35.8% |
| BUSCO completeness | 98.3% | 93.6% |

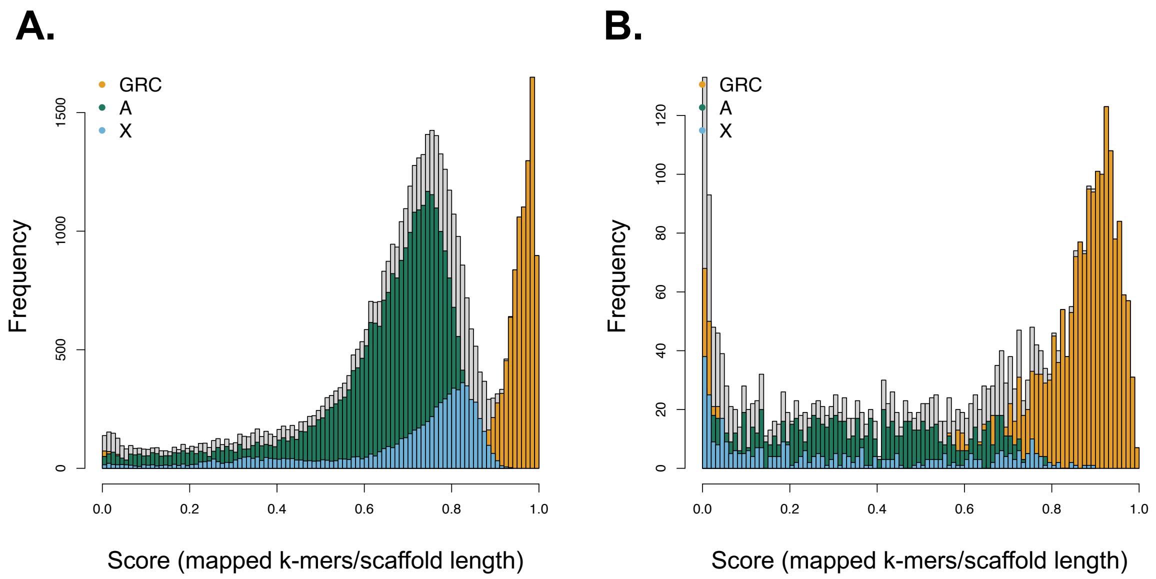


**Supplementary Fig 2. Distributions of scores used in the k-mer identification technique. A.** Histogram of k-mer assignment scores for scaffolds of each chromosome type in the short-read assembly used throughout the manuscript. The score is defined as the maximum number of k-mers with an exact match to the scaffold from the chromosome with the majority of k-mers matching the scaffold, divided by the scaffold length. For GRC scaffolds (orange), we only assigned scaffolds with a score higher than 0.8 as GRC scaffolds, while for autosomal and X chromosome scaffolds (green and blue respectively) we assigned scaffolds with a score higher than 0.4. **B.** Histogram of k-mer assignment scores in long read assembly (See **Supplementary Methods**). The scores are substantially lower, especially for differentiating autosomes and the X chromosome. This assembly was used only for anchoring GRC genes in longer blocks for the collinearity analysis.

**Supplementary Table 2**. Size and proportion of reference genome [16] anchored to specific chromosomes in *B. coprophila* and number of GRC paralogs and number of GRC collinear blocks anchored to each chromosome in the reference assembly.

| | A-II | A-III | A-IV | X |
|---|---|---|---|---|
| Size (Mb) | 48-62 | 66-71 | 88-94 | 48-62 |
| Proportion anchored | 20-46% | 8-19% | 37-52% | 93-100% |
| GRC paralogs (number genes) | 128 | 7 | 108 | 119 |
| GRC collinear blocks | 14 | 1 | 12 | 12 |

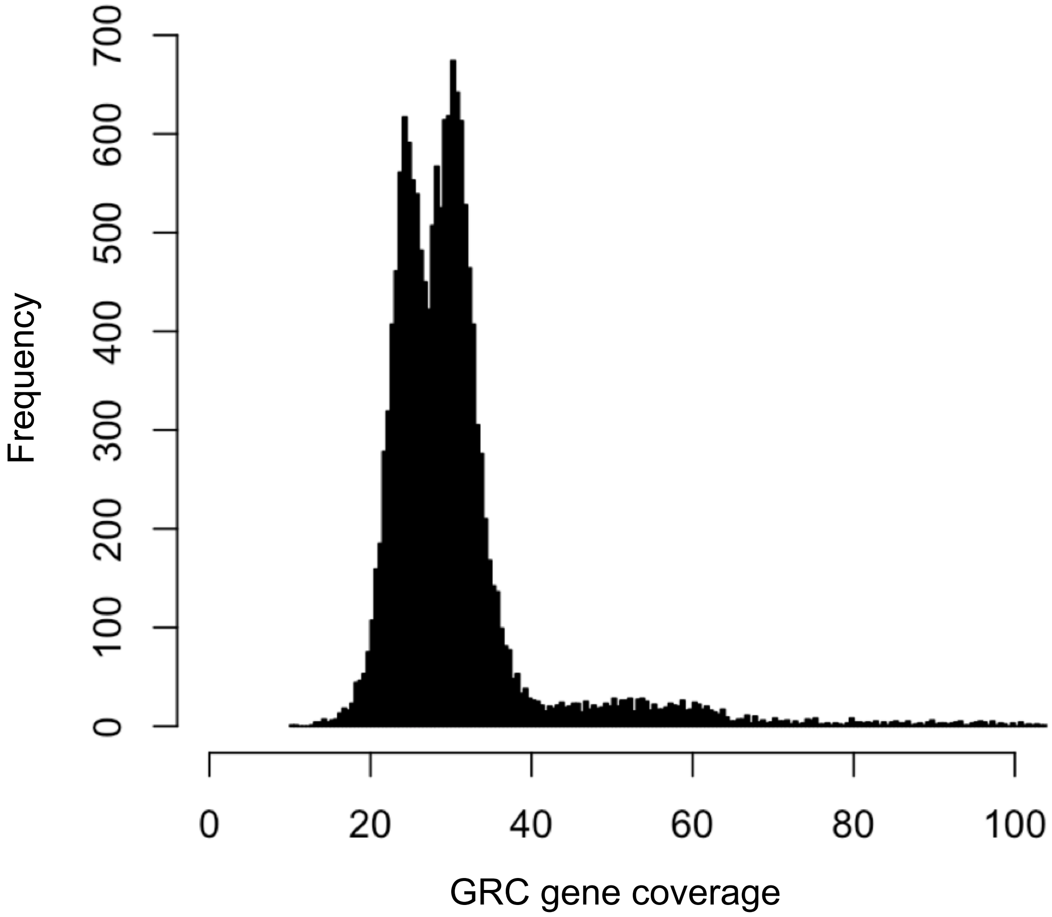


**Supplementary Fig 3. Histogram of the mean coverage of all GRC genes**. The coverage of GRC genes is bimodal, with one peak at 24.6x coverage and another at 30.3x coverage.

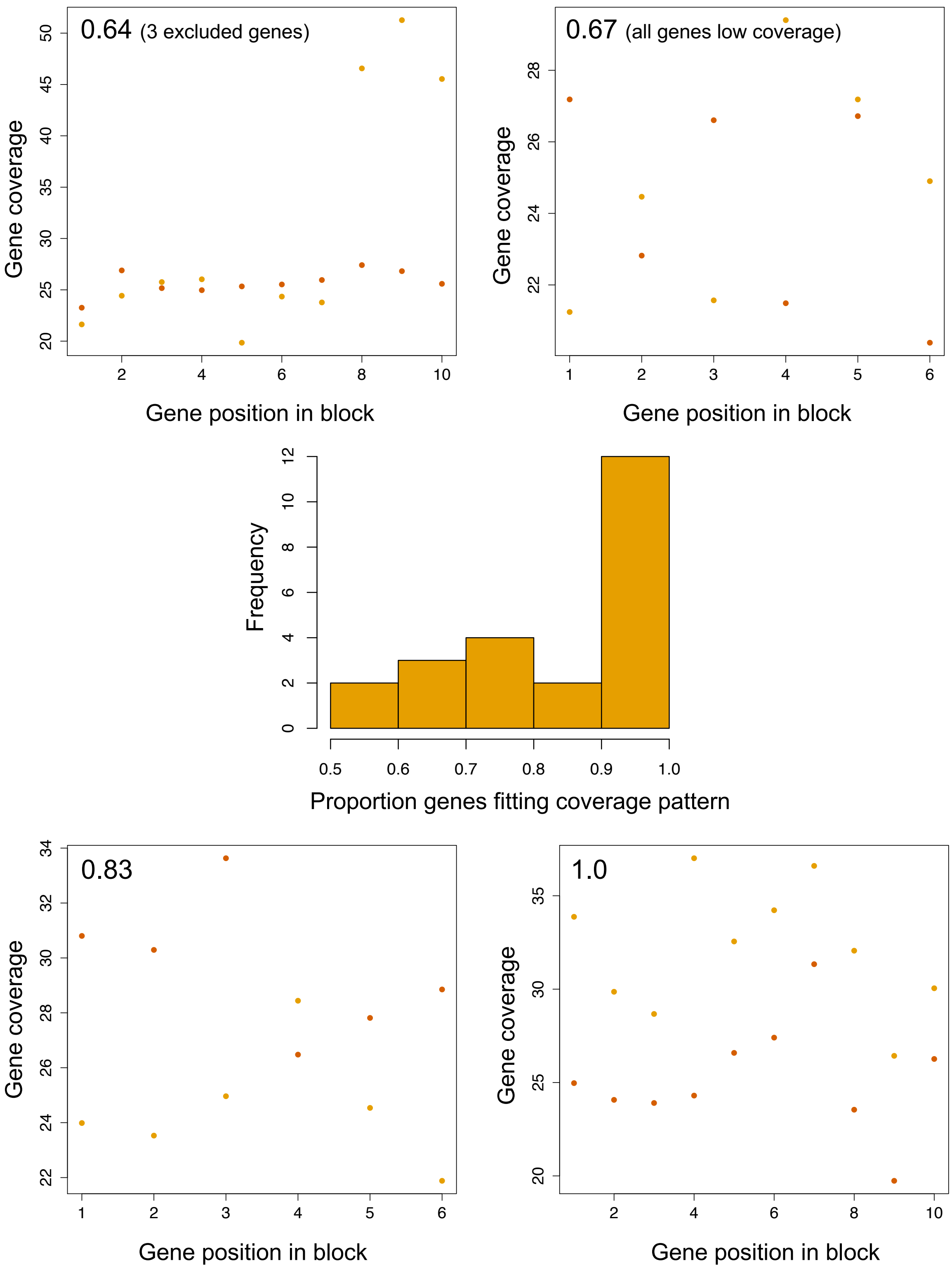


**Supplementary Fig 4. Coverage of genes in GRC-GRC collinear blocks.** The two blocks are coloured with different shades of orange. The histogram shows the proportion of genes in each block that fit the expected coverage pattern (i.e. one block having all genes with a higher coverage than the other). Only genes with a coverage less than 45x were considered.

Four blocks with different coverage patterns are shown with the score assessing how many paralogs fit the expected coverage pattern in the top corner of each plot. The plot in the top left corner has several genes with a higher coverage than expected (which were excluded), with the other paralogs in the block having inconsistent coverage patterns. The top right corner shows a case where all genes have coverage distributions which would fit in the lower coverage peak (in **Supplementary Fig 1**), and an inconsistent coverage pattern (likely because these blocks are both located on the same GRC), and the lower left corner shows a case where most paralogs fit the expected coverage pattern except one set of paralogs, which has an intermediate coverage level. Most blocks (13/23) show the same pattern as the block in the lower right corner, where all genes in one block have a higher coverage than the genes in the other block.

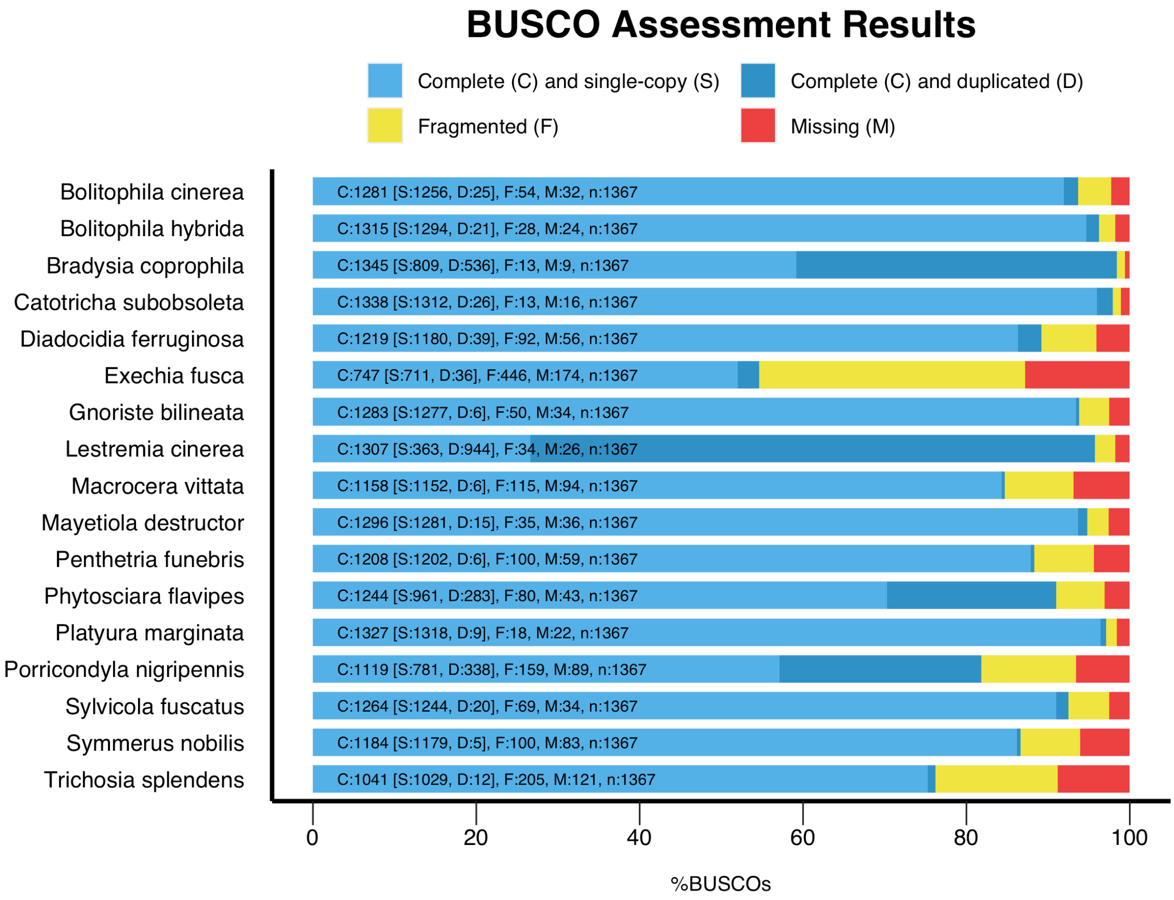


**Supplementary Fig 5. Summary of universal single-copy orthologs (BUSCO) results for all Dipteran species in phylogenetic analyses**. *Exechia fusca* was excluded from analyses as the proportion of complete BUSCOs was low (54%). In *Bradysia coprophila,* 39.2% of the insect BUSCO genes were duplicated.

**Supplementary Table 3. Genomic location of universal single-copy orthologs (BUSCO) in *Bradysia coprophila*.** BUSCO assessment was conducted with the insecta_odb10 database. The genomic location of genes was identified with both coverage and k-mer identification techniques. Categories indicated with * were used in phylogenetic analyses (**Fig 5A**) to determine the phylogenetic position of GRC genes, and individual gene trees were examined (**Fig 5C**) for the paralogs in the categories indicated with * and **.

| BUSCO type | Chromosome | Frequency | GRC related |
|---|---|---|---|
| Single-copy | A | 521 | No |
| | X | 182 | No |
| | GRC | 106 | Yes |
| Duplicated | A-GRC* | 291 | Yes |
| | X-GRC* | 81 | Yes |
| | GRC-GRC | 18 | Yes |
| | A-A | 6 | No |
| | A-X | 1 | No |
| Multi-copy | A-GRC-GRC** | 56 | Yes |
| | X-GRC-GRC** | 30 | Yes |
| | GRC-GRC-GRC | 3 | Yes |
| | A-A-GRC | 3 | Yes |
| | A-X-GRC | 2 | Yes |
| | A-GRC-GRC-GRC | 1 | Yes |
| | X-X-GRC-GRC-GRC | 1 | Yes |

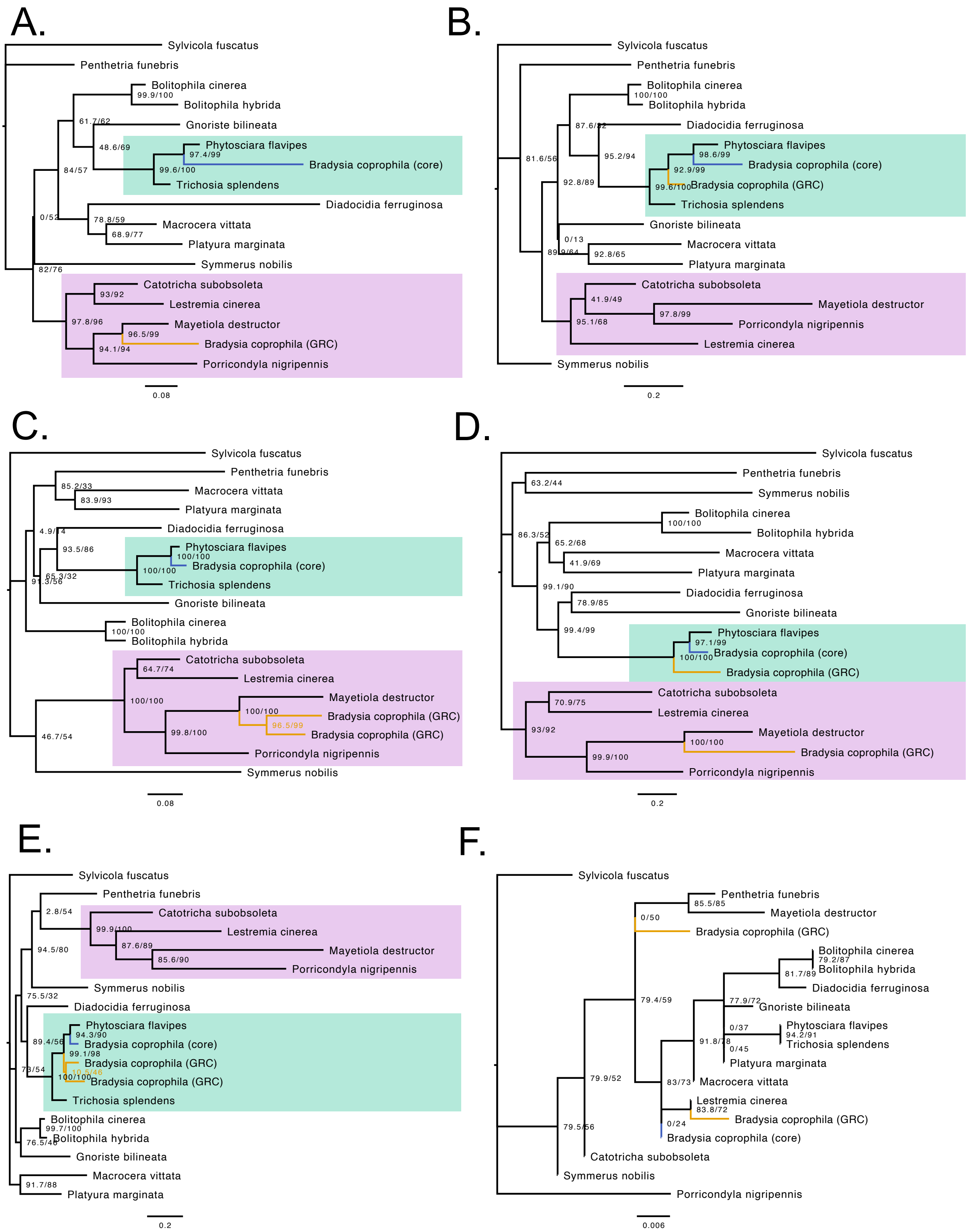


**Supplementary Fig 6. Examples of GRC gene trees with various topologies**. (A) with one GRC copy rooted in Cecidomyiidae (B) with one GRC copy rooted in Sciaridae; (C) with two GRC copies both in Cecidomyiidae (E) with two GRC copies, one in Cecidomyiidae and the other in Sciaridae (D) with two GRC copies both in Sciaridae. (F) GRCs unplaced (without significant nodes) or branching with a species from any other family. These six gene trees are representative examples of individual categories of gene tree topologies on **Fig 5B**.

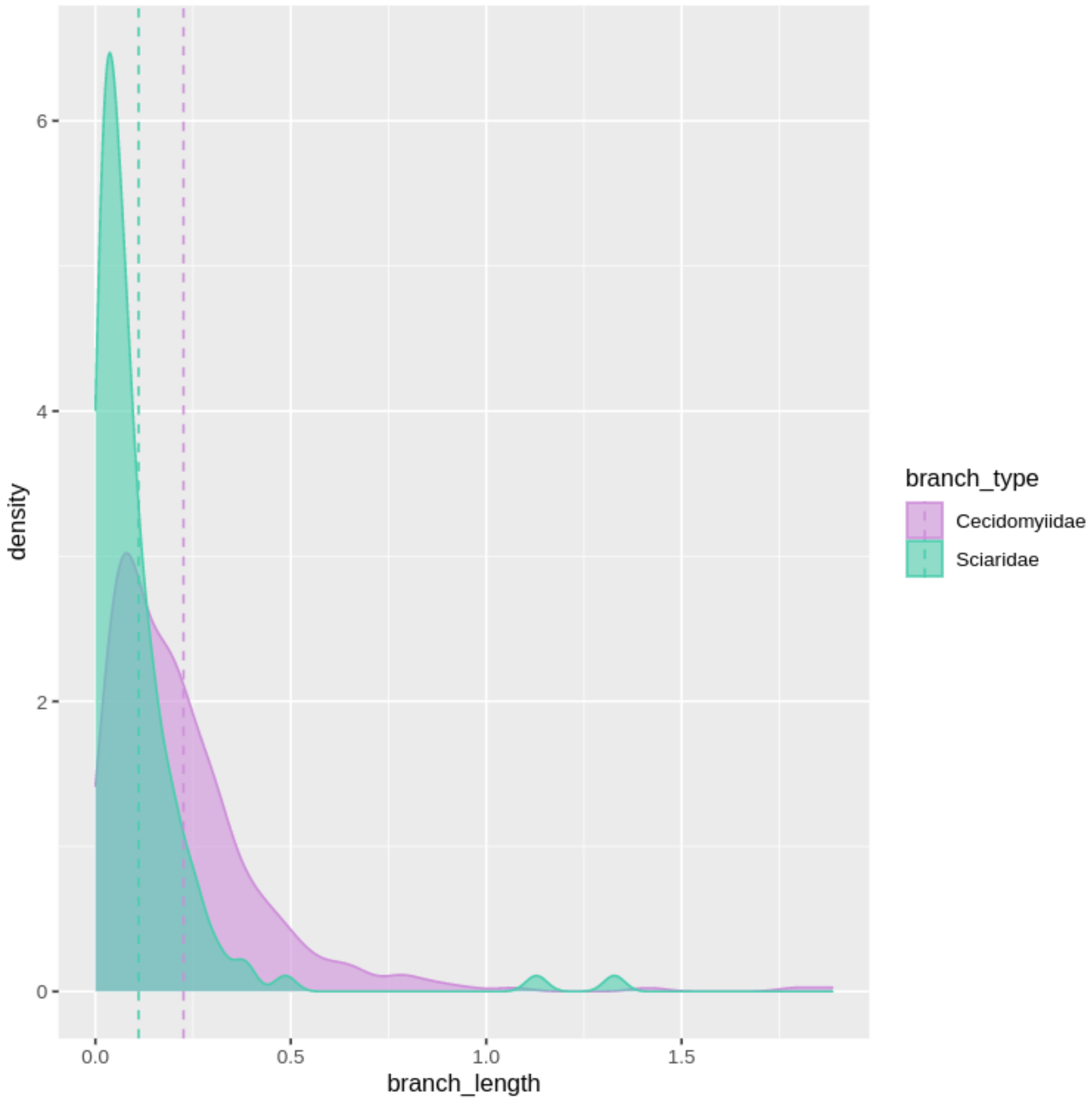


**Supplementary Fig 7. Terminal branch length distribution of GRC genes;** Branch length distribution of GRC copies of BUSCO genes plotted with respect to the phylogenetic position (at family level), means shown by dashed lines. Branch lengths of BUSCO genes on GRCs within Cecidomyiidae (violet) are significantly longer than branches found within Sciaridae (teal; p-value $< 0.0001$) suggesting the genes on GRCs found within Sciaridae might be due to gene duplications and translocations within Sciaridae after the GRCs were acquired.

**Supplementary Text 3: Approximate dating of the introgression of GRCs**

We used Baltic amber records to roughly estimate the date of the introgression of GRCs from Cecidomyiidae to Sciaridae. The records of the early diversification of the Sciaridae family found in the amber are dated to be ~44myo [17,18]. Supposedly, the common ancestor of the family is a bit older than that, therefore, we roughly estimate the common ancestor of Sciaridae to be 50 mya. We can use this estimate to time calibrate the phylogeny of 340 BUSCO genes (see **Fig 5**). Using this logic, the common ancestors of Sciaridae and Cecidomyiidae was 147 mya (calculated using sum of branches from Sciaridae backwards). The isolation of Sciaridae then happened ~31 my after the split with Cecidomyiidae. We hypothesise the introgression must have happened after that as no other Bibionomorpha families have GRCs nor show any other signs of hybridization with ancestors of Cecidomyiidae [19,20]. The hybridisation therefore probably happened on the Sciaridae branch before the diversification of the family, approximately 116 - 50 mya and between 31 - 97 my after the split of the original ancestors of the two families.

These calculations must be taken with a large grain of salt, as the substitution rates change over time and our calibration is relatively crude as it uses only one reference point. However, it suggests that hybridisation of extremely divergent species (31 - 97 my) can have important evolutionary consequences. This is not the first record of viable hybrids of two extremely diverged animals. Recently a successful hybridization of Russian Sturgeon and American Paddlefish was accomplished [21]. However, this was a lab-generated hybrid, not a result of mating in the wild. Other hybridisation events of diverged animals that resulted in gene flow are found in *Nasonia wasps* [22], sea squirt [23], and burrowing frogs [24]. The B chromosome in *Nasonia* wasps is thought to have arisen through hybridization with a *Trichomalopsis* wasp, the two lineages are estimated to diverge for 2.6 my [25]. The B chromosome also have a substantial effect on *Nasonia* wasps, as it affects sex determination in individuals that carry it. The sea squirt the gene flow appeared after secondary contact of more than 3 my of divergence of the two lineages [23], which is already an upper boundary of known systems with ongoing gene flow. Gene flow between more remote lineages, as in burrowing frogs, seems to be facilitated by polyploidisation [24]. Similar case is found in *Arabidopsis lyrata* and *A. arenosa* species complex. Both those species have diploid and tetraploid forms and while the diploid variants are fully reproductively isolated, the tetraploid variants form viable hybrids generating an indirect route for gene flow between these ~20my diverged lineages [26]. It appears that hybridization events of extremely diverged species are always associated with polyploidy. Isolation of the two genomic copies in separate instances increases the stability as the recombination does not break up already-working of within-subgenome co-adapted genes. It seems likely, given that the size of the GRCs in *B. coprophila* is comparable to the size of the entire Cecidomyiid genome (the genome size of *Mayetiola destructor,* for example, is 158Mb; [27]), that the originally introgressed GRCs in the common ancestor of Sciaridae carried a full genomic copy of the ancestral Cecidomyiidae genome. We speculate that GRCs originated in Sciaridae through introgression of the full cecidomyiid genome. The introgression might have directly resulted in GRCs or was followed by restriction of the introgressed genome to the germline. Sciaridae species therefore represent a rare case of germ-line specific polyploids.

21. Káldy J, Mozsár A, Fazekas G, Farkas M, Fazekas DL, Fazekas GL, et al. Hybridization of russian sturgeon (Acipenser gueldenstaedtii, Brandt and Ratzeberg,

1833) and american paddlefish (Polyodon spathula, Walbaum 1792) and evaluation of their progeny. Genes (Basel). 2020;11: 1–17. doi:10.3390/genes11070753

27. Stuart JJ, Chen M, Harris MO. Hessian fly. Publ from USDA-ARS / UNL Fac US. 2008.
